# Direct targeting of the WAVE-regulatory complex and actin cytoskeleton by the *Anaplasma phagocytophilum* effector, AnkA

**DOI:** 10.64898/2026.07.31.742066

**Authors:** Hannah L. Burge, Shashi P. Singh, Jianyang Wang, Suchitra Pradhan, Katharine M. Wood, Joshua W. Lee, Rob Barringer, Silvia A. Synowsky, Sally L. Shirran, Patrick J. Moynihan, Andrew L. Lovering, Laura M. Machesky, Mark A. Jepson, J. Stephen Dumler, Ian T. Cadby

## Abstract

The obligate intracellular tick-borne pathogen *Anaplasma phagocytophilum* is unusual in its tropism for neutrophils, within which it survives and replicates. Central to colonization of this hostile niche are secreted effector proteins. One such effector protein, AnkA, interacts with multiple host ligands in the cytoplasm and nucleus, yet how these interactions promote infection remains unclear. We discovered that AnkA directly binds both the WAVE Regulatory Complex (WRC), a signal-integration hub regulating branched actin networks, and monomeric and filamentous actin. AnkA uses two forms of molecular mimicry: binding to the Abi1 component of the WRC via a WRC-Recruiting Abi1-binding Peptide (WRAP), and binding actin via a WH2-like Actin Binding Motif (ABM). These interactions enable AnkA to exert multi-factorial influence on the actin cytoskeleton and signaling pathways converging on this structure. AnkA and the WRC co-localise during bacterial internalization and late stages of infection, suggesting roles in both host-cell colonization and bacterial dissemination. Our work reveals AnkA as the first example of a bacterial effector that directly targets the WRC. The co-occurrence of WRAP- and ABM-like motifs in other bacterial effector proteins suggests that this mechanism of actin cytoskeleton subversion may represent a conserved strategy used by other obligate intracellular bacteria.

## Introduction

The host cytoskeleton is a common target of many infectious microbes, being hijacked to facilitate a great many aspects of infection [1–3]. Obligate intracellular bacteria, being wholly reliant on host cells for replication, are constitutively exposed to the host cell cytoskeleton because of their lifestyle. However, owing to challenges to their study, such as comparatively poorly developed genetic approaches, our understanding of how most obligate bacteria interface with and manipulate the host cytoskeleton is limited [1].

*Anaplasma* is a genus of obligate intracellular bacteria that infect a wide range of vertebrates in addition to their primary vectors, ticks. Diseases caused by *Anaplasma* have major impacts on global livestock industries and are amongst the most prevalent tick-borne diseases worldwide [4–6]. *Anaplasma phagocytophilum* is a zoonotic pathogen of humans, livestock, and domestic animals, and the only bacterium known to specialize in infecting and replicating within host neutrophils. This is an unusual niche since neutrophils are short-lived and potently equipped to kill microbes [7–9].

In the host cell, *A. phagocytophilum* resides and replicates within a host-derived membrane-bound vacuole (the ApV) which has similarities to early autophagosomes [8, 10]. To facilitate this lifestyle, *A. phagocytophilum* employs a type IV secretion system which enables the delivery of bacterial effector proteins directly into the host cell [11, 12]. Only a handful of *A. phagocytophilum effectors* have been experimentally characterized and in most cases, the mechanisms underpinning their functions are unknown [12–18].

The AnkA effector was first identified as a ∼160 kDa serological marker of *A. phagocytophilum* infection, indicating that it is expressed within the vertebrate host [13, 19]. Named due to the presence of multiple ankyrin repeats in its protein sequence, AnkA is conserved throughout genus *Anaplasma* and is essential for survival in mammalian cell culture [20]. AnkA fulfills diverse functions. Within the host cytoplasm, AnkA is tyrosine phosphorylated by host kinases Src and Abl on EPIYA motifs, pathogenic protein motifs found in a range of structurally unrelated bacterial effectors which promote binding with SH2-domain containing proteins such as SHP-1 [13, 21]. AnkA also interacts with the host adapter protein Abi1 and it is thought that this interaction enables the formation of a tripartite complex comprised of AnkA, Abi1, and Abl kinase [13]. Most recently, AnkA has been implicated in influencing the actin cytoskeleton by binding actin, gelsolin, and α-actinin-4 [22]. AnkA is also translocated to the nucleus wherein it binds to DNA and the nuclear protein HDAC1, influencing gene expression and epigenetics [23–26]. Despite it being apparent that AnkA interacts with many host targets, the consequences of these interactions on host biology are incompletely defined, and it is likely that additional AnkA-targets are yet to be identified [23].

In this study, we sought to further our understanding of the role of AnkA in host subversion by using unbiased approaches to identify mammalian host cell AnkA interaction partners and define the influence of AnkA on their biochemistry. In doing so, we discover new roles for AnkA in the direct and indirect manipulation of host actin via molecular mimicry.

## Results

### AnkA interacts with the WAVE Regulatory Complex and actin in mammalian cells in a domain dependent manner

We sought to identify AnkA interaction partners via an immunoprecipitation and proteomics strategy. Polyclonal anti-AnkA antibodies were used to immunoprecipitate AnkA from lysates generated from infected and uninfected HL-60 cells, a human leukemic cell-line with neutrophil-like properties used for laboratory culture of *A. phagocytophilum*. Candidate AnkA interaction partners were identified by Label-free liquid chromatography-mass spectrometry/mass spectrometry (LC-MS-MS). AnkA and numerous other proteins were enriched from infected cell lysates (**Fig. 1A, Supplementary xls file 1**). Of particular interest amongst this list was Abi1, a mammalian host protein previously reported to bind AnkA [13], and four additional proteins; CYFIP1/2, NCKAP1L, WAVE1/2, and BRK1; which associate with Abi1 and form the pentameric WAVE Regulatory Complex (WRC, **Fig. 1A**). Actin, whilst present in both samples, was also enriched by anti-AnkA from infected cell lysates.

**Figure 1.**
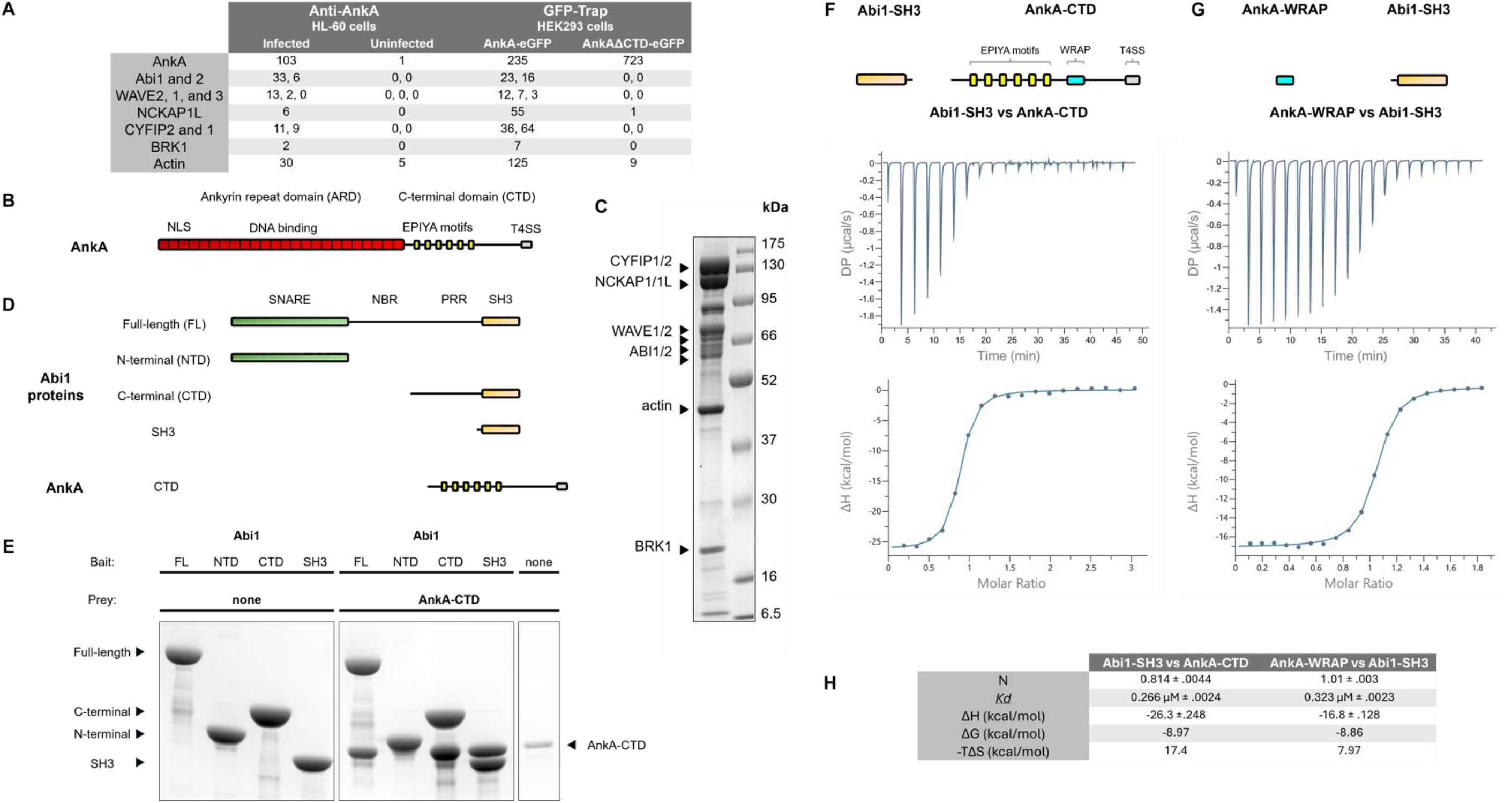
AnkA binds to the WRC via the SH3 domain of Abi1. A. LC-MS-MS peptide counts of candidate AnkA interaction partners identified by immunoprecipitation of AnkA *from A. phagocytophilum* infected HL-60 cell lysates (first two columns) and immunoprecipitation of AnkA-GFP and AnkAΔCTD-GFP from transfected HEK293 cells. B. Schematic of the full-length AnkA protein. Characterised regions are labelled: nuclear localisation signal (NLS), DNA binding domain, EPIYA motifs, and Type IV secretion signal (T4SS). C. SDS-PAGE gel of proteins isolated from porcine brain lysate using AnkA-CTD beads. The anticipated molecular weights of the individual WRC units, marked by arrows, are Cyfip1/2 (145 kDa), Nckap1 (129 kDa), WAVE1/2 (54-62 kDa), Abi1 (52-59 kDa), BRK1 (11 kDa). Wave, Abi and Brk units are subject to multiple post-translational modifications leading them to run at different apparent molecular weights. D. Schematic of Abi1 proteins and the AnkA-CTD protein used in pull-down assays. Domains are labelled: SNARE, Nap1 binding region (NBR), proline rich region (PRR), and SH3. E. Coomassie-blue-stained SDS-PAGE gel of pull-down experiments. MBP-tagged Abi1 proteins were bound to amylose beads and used as bait with AnkA-CTD protein as prey. F. Representative isothermal calorimetry isotherm for the binding of Abi1-SH3 to AnkA-CTD and integrated heats fitted to a single-binding site model. A schematic of the proteins used is shown above the plots. G. Representative isotherm for the binding of an AnkA-derived peptide, AnkA-WRAP, to Abi1-SH3 and integrated heats fitted to a single-binding site model. Results are representative of two independent experiments. H. Dissociation constants and thermodynamic parameters from ITC experiments.

As a complementary approach, we expressed AnkA-eGFP and eGFP only fusion proteins in the human cell line HEK293 by transient transfection and then used GFP-Trap to immunoprecipitate AnkA-eGFP and eGFP from cell lysates. LC-MS-MS analysis demonstrated that the WRC also co-immunoprecipitated with ectopically expressed AnkA-eGFP but not with eGFP alone (**see Supplementary xls file 2**). Notably, WRC proteins were the only candidate AnkA-interaction partners common to both IP enrichment strategies.

Having co-immunoprecipitated with AnkA using two distinct capture methods, we deduced that the WRC was likely a bona fide interaction partner of AnkA. The WRC is a WASP-family protein complex that functions as a signalling hub, regulating the formation of branched actin networks via the downstream Arp2/3 complex [27]. In its basal state, the WRC is auto inhibited but signalling proteins (Rac and Arf) can activate the complex, leading to the release of the WAVE VCA domain which in turn interacts with actin and Arp2/3, activating the latter [28–31]. This activity drives key functions including cell migration and phagocytosis in multiple cell types, including neutrophils [32].

The AnkA protein is comprised of two domains: an N-terminal domain of ∼900 amino acids comprised mainly of ankyrin or ankyrin-like repeats and which we term the ankyrin repeat domain (ARD), and a C-terminal domain of ∼350 amino acids that is predicted to be mainly disordered peptide and which we term the C-terminal domain (CTD, **Fig. 1B**). *A. phagocytophilum* is extremely challenging to manipulate genetically so, to determine which domain of AnkA binds the WRC, we expressed AnkA-eGFP with the CTD deleted (AnkAΔCTD-eGFP) in HEK293 cells and revisited our GFP-Trap based immunoprecipitation and proteomics strategy, using cells expressing AnkA-eGFP as a control. Deletion of the AnkA CTD resulted in loss of WRC and actin binding by AnkA (**Fig. 1A, also see Supplementary xls file 3**).

To confirm the role of the CTD in AnkA binding to the WRC, we generated AnkA-CTD covalently coupled beads and used these for a pull-down from porcine brain lysates (**Fig. 1C)**. We were able to capture all five components of the WRC and actin by this strategy (**Fig. 1C**). Across all three experiments, the WRC proteins were the only mammalian proteins specifically and robustly enriched by all IP strategies. We did not see any enrichment of previously reported interaction partners, Abl kinase and α-actinin-4, and gelsolin was only enriched in one of the three datasets.

Taken together, these results demonstrate that AnkA interacts with the WRC and actin, and that these interactions are mainly dependent upon the CTD of AnkA. These results also indicate a function for AnkA in exerting influence on the actin cytoskeleton via interaction with the WRC and direct binding to actin.

### AnkA directly interacts with the WRC via the SH3 domain of Abi1

The interaction between AnkA and Abi1 was first identified in a yeast-two-hybrid screen [13]. Since yeast genomes do not contain genes encoding WRC proteins [27] we therefore posited that Abi1 is involved in mediating the interaction between AnkA and the WRC.

The Abi1 protein is comprised of multiple domains (**Fig. 1D**). The N-terminus, comprising the SNARE and Nap1-binding fragment (NBR) domains, interacts with other WRC proteins, whereas the C-terminal proline-rich region (PRR), and Src homology 3 (SH3) domains do not [33]. As such, the C-terminal domains of Abi1 are free to bind other SH3-containing proteins (via the Abi1-PRR) and/or to other proline rich proteins (via the Abi1-SH3), recruiting signaling proteins to WRC and influencing its activity and cellular localisation [34, 35].

To define which domain AnkA interacted with, we used domain-deleted Abi1 recombinant proteins in pull-down assays with purified AnkA-CTD protein (**Fig. 1D & E)**. Abi1 proteins were bound to amylose beads and used as bait in pull-down assays with purified recombinant AnkA-CTD protein as prey (**Fig. 1E**). AnkA-CTD bound to Abi1 proteins that included the SH3 domain. Whilst these results do not preclude the possibility that AnkA forms other contacts with the WRC, they do indicate that the SH3 domain of Abi1 alone is sufficient to mediate interaction with the AnkA-CTD.

We next used isothermal calorimetry (ITC) to define the stoichiometry and thermodynamics of the interaction between recombinant AnkA-CTD and Abi1-SH3 proteins (both purified free of affinity tags, **Fig. 1F**). Sequential injections of Abi1-SH3 into AnkA-CTD resulted in measurable heat changes and proceeded to saturation (**Fig. 1F)**. The resulting isotherm was fitted to a single binding site model and thermodynamic parameters associated with the AnkA-CTD:Abi1-SH3 interaction were extracted (**Fig. 1H).** Abi1-SH3 binds to AnkA-CTD with a *Kd* of 266 nM and a binding stoichiometry near 1:1, confirming direct and relatively high affinity between these protein domains. AnkA-CTD binding to Abi1-SH3 domains is primarily enthalpy driven (ΔH −26.3) and proceeded with a positive -TΔS (17.4), indicating a large change from disorder to order in the binding of AnkA-CTD (which is predicted to be mostly disordered) with Abi1-SH3.

SH3 domains are found in a range of signaling proteins and generally bind to proline rich sequences, particularly PXXP motifs where X denotes any amino acid [36]. Using the Eukaryotic Linear Motif (ELM) server [37], we identified a potential SH3 interaction motif with the sequence KVKPQVP in the AnkA-CTD. A peptide of 21 amino acids, based on AnkA and centered on this candidate SH3 interaction motif, was synthesized and used in ITC experiments. Titration of the peptide into Abi1-SH3 resulted in heats of binding and isotherms very similar to those obtained with AnkA-CTD and Abi1-SH3 (**Fig. 1G & H)**, indicating the peptide represents the core Abi1-SH3 binding residues of AnkA.

These results demonstrate that AnkA interacts directly with the SH3 domain of Abi1, a component of the WRC, via a discrete motif within the AnkA-CTD. Accordingly, we term this motif the **W**RC-**R**ecruiting **A**bi1 binding **P**eptide, or WRAP (**Fig 1F & G)**.

### The structure of an AnkA-WRAP derived peptide in complex with Abi1-SH3

To identify the specific residues that mediate the interaction and glean clues to the likely biochemical consequences that AnkA binding would have on Abi1 and, by extension, the WRC, we determined the crystal structure of the AnkA-WRAP peptide bound to Abi1-SH3 domain. The AnkA-WRAP:Abi1-SH3 complex crystallized in space group *C* 1 2 1 and was solved to a resolution of 1.69 Å **(Fig. 2A, Supplementary table 1)**. The asymmetric unit comprises two copies of the protein complex and the majority of the polypeptide chains used in the crystallography experiments were resolved (AnkA residues 1096-1116 and Abi1 residues 429-503). The Abi1-SH3 domain is comprised of a five-stranded beta-barrel fold with a short 3_10_ helix separating strands β4 and β5, typical of the SH3 family [36]. Residues W483, P496, Y499, Y455 form a shallow groove for PXXP motif binding. Two loops at the end of the PXXP binding groove, one between β1 and β2 (termed the RT-loop) and one between β3 and β4 (termed the n-Src loop), form a cleft called the specificity site and which confers additional binding selectivity. SH3 binding proteins are classified according to their orientation of binding [38].

**Figure 2:**
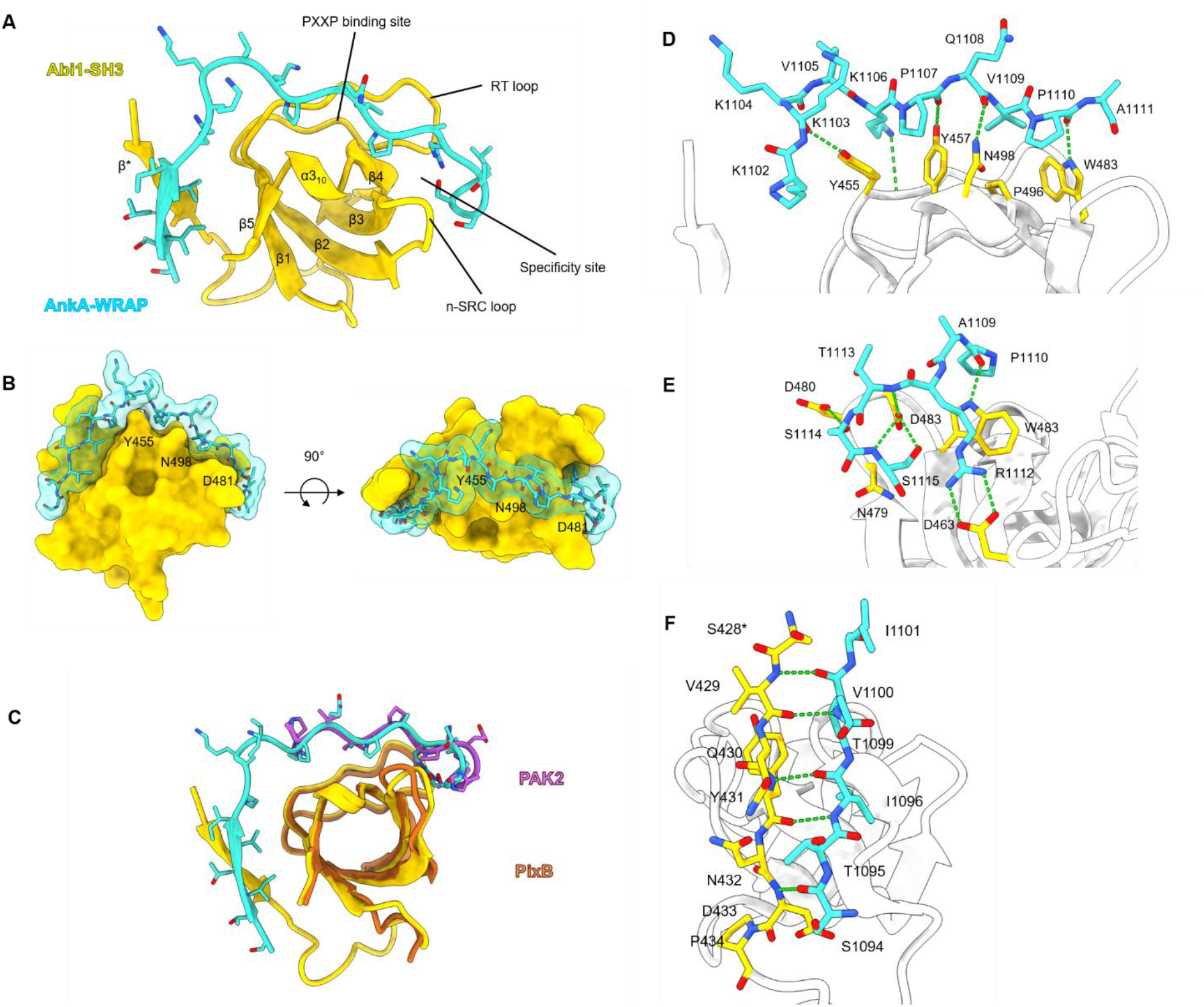
Crystal structure of Abi1-SH3 in complex with the AnkA-derived peptide, AnkA-WRAP, solved to 1.69 Å. A. Cartoon representation of AnkA-WRAP in complex with Abi1-SH3. Defined structural elements of the SH3 domain are labelled. The protein complex comprises AnkA residues 1096-1116 and Abi1 residues 429-503. B. Surface representation of the AnkA-WRAP:Abi1-SH3 complex. AnkA-WRAP wraps around Abi1-SH3, binding via shape complementarity. C AnkA-WRAP:Abi1-SH3 superimposed onto the structure of PixB in complex with a PAK2 derived peptide. D. Binding of Ank-WRAP to the PXXP binding site of Abi1. E. Interactions between AnkA-WRAP and the specificity site of Abi1-SH3. F. Non-canonical interactions between AnkA-WRAP and β* of Abi1-SH3. *Residue S428 is a remnant of the TEV cleavage site used in the production of recombinant Abi1-SH3 and is equivalent to A428 of Abi1.

The AnkA-WRAP peptide binds to the Abi1-SH3 PXXP and specificity sites in the N- to C-terminus direction, consistent with being a Class II SH3-interaction partner. Consistent with this, the AnkA sequence, PQVPAR, matches the consensus (PXXPXR) for a Class II SH3 binding peptide. Binding is largely mediated by shape complementarity (**Fig. 2B).** The AnkA-WRAP motif mimics endogenous SH3 binding proteins but has an extended interface and forms non-canonical contacts outside of the PXXP and specificity sites (**Fig. 2C)**.

AnkA forms mixed interactions with the Abi1-SH3 domain. At the PXXP binding site, AnkA residues P1107, V1109 and P1110 interdigitate with Abi1 aromatic residues Y455, Y457, W483 and Y499 **(Fig. 2D)**. At the specificity site, AnkA residue R1112 forms a salt-bridge with Abi1 residue D463 and stacks with Abi1 W483. Also at this site, Abi1 D480 and D481 form a network of hydrogen bonds with the AnkA backbone and side chains of T1113, S1114 and S1115 (**Fig. 2E)**. In addition to these canonical SH3 interactions, AnkA residues 1096-1101 and the Abi1-SH3 β* residues form backbone interactions to form two antiparallel inter-chain β strands (**Fig. 2F**). Additional hydrophobic contacts are made between AnkA I1098 and V1100, and Abi1 I454 and I503 of β1 and β5, respectively. We posit that the disorder to order transition measured by ITC on AnkA-WRAP:Abi1-SH3 binding is partly reflective of AnkA forming β-interactions with the Abi1-SH3 β*.

In summary, the 21 amino acids of the AnkA-WRAP peptide wraps around the Abi1-SH3 domain, forming canonical and non-canonical interactions, effectively covering 180° of the SH3 surface on this plane (**Fig 2B)**. The extensive interface suggests relatively high binding specificity or preferential binding between AnkA-WRAP and Abi1-SH3, a suggestion supported by the absence of other SH3 domain containing proteins in our IP-proteomics datasets. AnkA mimics endogenous SH3-binding proteins and so its binding to Abi1-SH3 and the binding of Abi1-SH3 to other endogenous SH3 binding partners are expected to be mutually exclusive (**Fig. 2C**). In line with this, we recorded no peptide counts for known Abi1 SH3 binding partners, such as Abl kinase or Caskin2 in our IP datasets [39, 40].

### AnkA localises to lamellipodia in model expression systems

The WRC plays a central role in directed cell motility as an activator of the Arp2/3 complex in lamellipodia [29] so we next asked whether AnkA would localise to lamellipodia when heterologously expressed in mammalian cells. AnkA-eGFP or eGFP alone were expressed in the mammalian cell line B16F1 which were then imaged to observe lamellipodia formation and cell migration (**Fig. 3**). B16F1 cells, whilst not relevant to *A. phagocytophilum* infection biology, are a favourable cell line for the study of cell migration and WRC functions due to their relative ease of transfection compared to HL-60 cells and for forming large lamellipodia. B16F1 cells expressing either AnkA-eGFP or GFP were able to form lamellipodia and migrate (**Fig. 3A & Supplementary video 1**). Whilst B16F1 cells expressing eGFP-only showed GFP fluorescence in the cytoplasm, AnkA-eGFP was recruited at the leading edge of lamellipodia in migrating cells (**Fig. 3A & Supplementary video 1).** Similar experiments with fluorescent mCherry-tagged Abi1 yielded a localisation pattern matching that of AnkA-eGFP (**Fig. 3B & Supplementary video 2).** When AnkA-eGFP and Abi1-mCherry were co-expressed in B16F1 cells both proteins co-localised at the lamellipodia edge (**Fig. 3C & D**).

**Figure 3:**
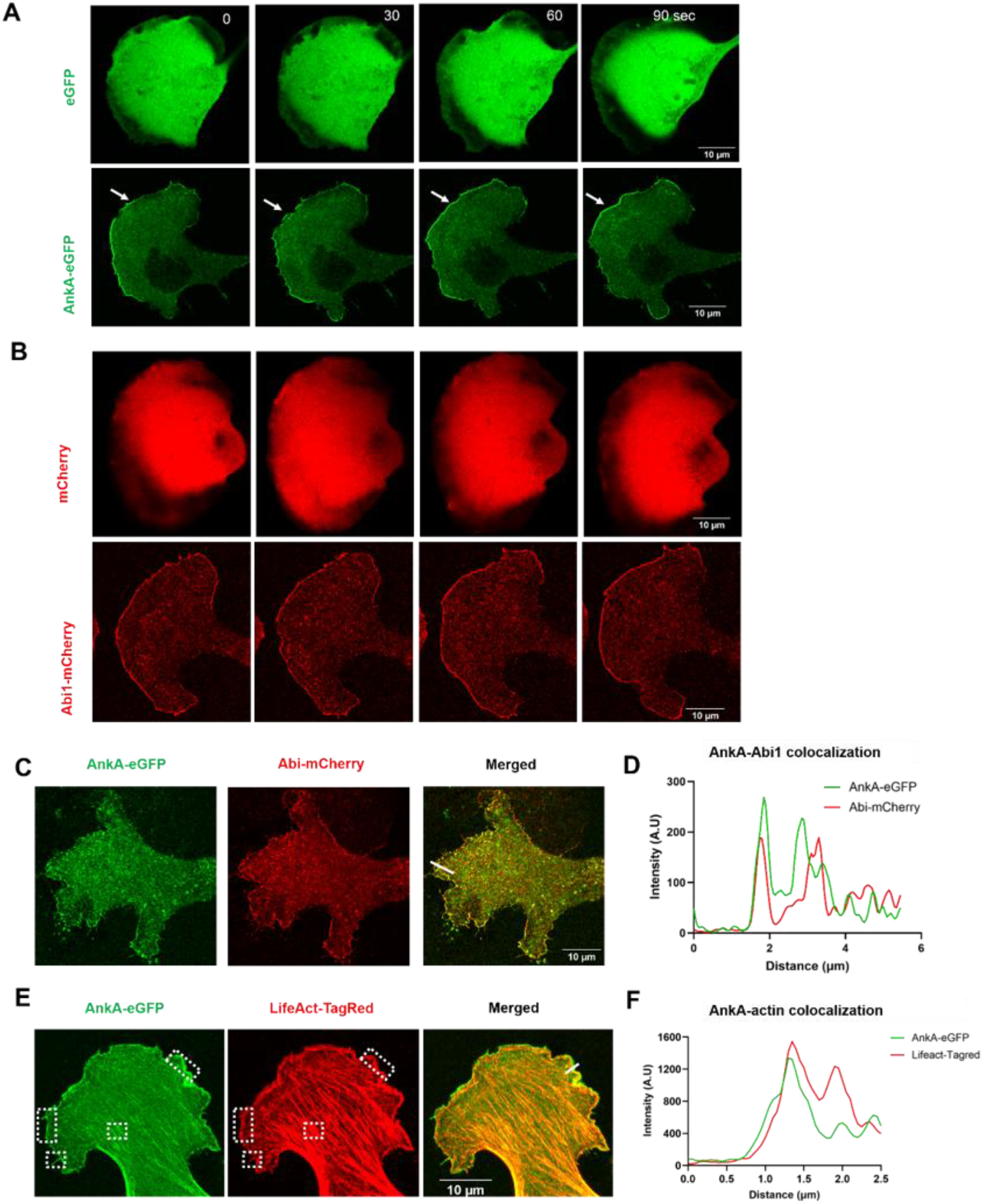
AnkA-eGFP localises to the lamellipodia with Abi1-mCherry and with actin rich structure in B16F1 cells. A. Cells were transfected with: pEGFP-C1 or AnkA-eGFP expression plasmids and imaged by live cell confocal microscopy. Recruitment of eGFP in the cytosol and AnkA-eGFP at the edge of the lamellipodia (white arrow). B. Cells were transfected with pMCherry-C1 or Abi1-mCherry expression plasmids and imaged by live cell confocal microscopy. Recruitment of mCherry in the cytosol and Abi1-mCherry at the edge of the lamellipodia (white arrow). C. Cells were co-transfected with AnkA-eGFP or Abi1-mCherry expression plasmids and imaged by live cell confocal microscopy. Co-recruitment of the AnkA-eGFP and Abi1-mCherry at the lamellipodia edge. D. Line scan of the lamellipodia edge (white line) indicating fluorescent intensities of the AnkA-eGFP and Abi1-mCherry corecruitment. E. Confocal microscopy image of B16F1 cell expressing AnkA-eGFP and LifeAct-TagRed. Dotted rectangular boxes showing recruitment of the two proteins. F. Line scan of the lamellipodia edge (white line) indicating fluorescent intensities of the AnkA-eGFP and LifeAct-TagRed co-recruitment at the leading edge and actin stress fibers.

These results indicate that ectopically expressed AnkA localises to the lamellipodia of migrating cells in a manner similar to Abi1 and that ectopically expressed AnkA and Abi1 proteins can be co-recruited to the lamellipodium. This localisation pattern of AnkA-eGFP is consistent with its role as a WRC interacting protein.

### AnkA localises to actin filaments and filopodia in a model system

Having defined the interaction between AnkA and the Abi1 component of the WRC we next sought to determine whether AnkA also exerts influence downstream of the WRC, on Arp2/3 and/or actin. We began by investigating whether AnkA co-localises with actin structures when heterologously expressed in mammalian cells. AnkA-eGFP and LifeAct-TagRed, which stains F-actin structures in cells, were co-expressed in B16F1 and imaged by confocal microscopy. Actin-rich structures including actin stress-fibres and filopodia were visible in LifeAct stained cells (**Fig. 3E & F).** AnkA-eGFP localised to stress-fibres and filopodia, indicating an association with F-actin structures. AnkA-eGFP also localised to the lamellipodium with a tight band present beyond the actin mesh, indicating that not all AnkA-eGFP at the lamellipodium is associated with F-actin.

### The AnkA-CTD directly interacts with both F- and G-actin

IP-proteomics and pull-down experiments demonstrated that the AnkA-CTD domain binds actin (**Fig. 1A**). Using the ELM server we identified a candidate WH2 actin binding motif downstream of the AnkA-WRC motif at AnkA residues 1133-1150 (sequence SSSFAAELQAQRGKLRPV). Directed by this prediction, we generated a series of recombinant MBP-tagged AnkA-CTD proteins with sections of the predicted disordered CTD deleted and used these in a series of F-actin pull-down assays to assess their ability to bind to actin filaments (**Fig. 4A).** SDS-PAGE analysis of actin pellets revealed that the AnkA-CTD co-pelleted with F-actin whereas an MBP only control did not (**Fig. 4B).** Deletion of AnkA residues 1135-1147, corresponding to the predicted actin binding motif, from any construct led to a decrease in actin binding, confirming that these residues impart F-actin binding. Accordingly, we have named this region of AnkA the actin binding motif or AnkA-ABM.

**Figure 4:**
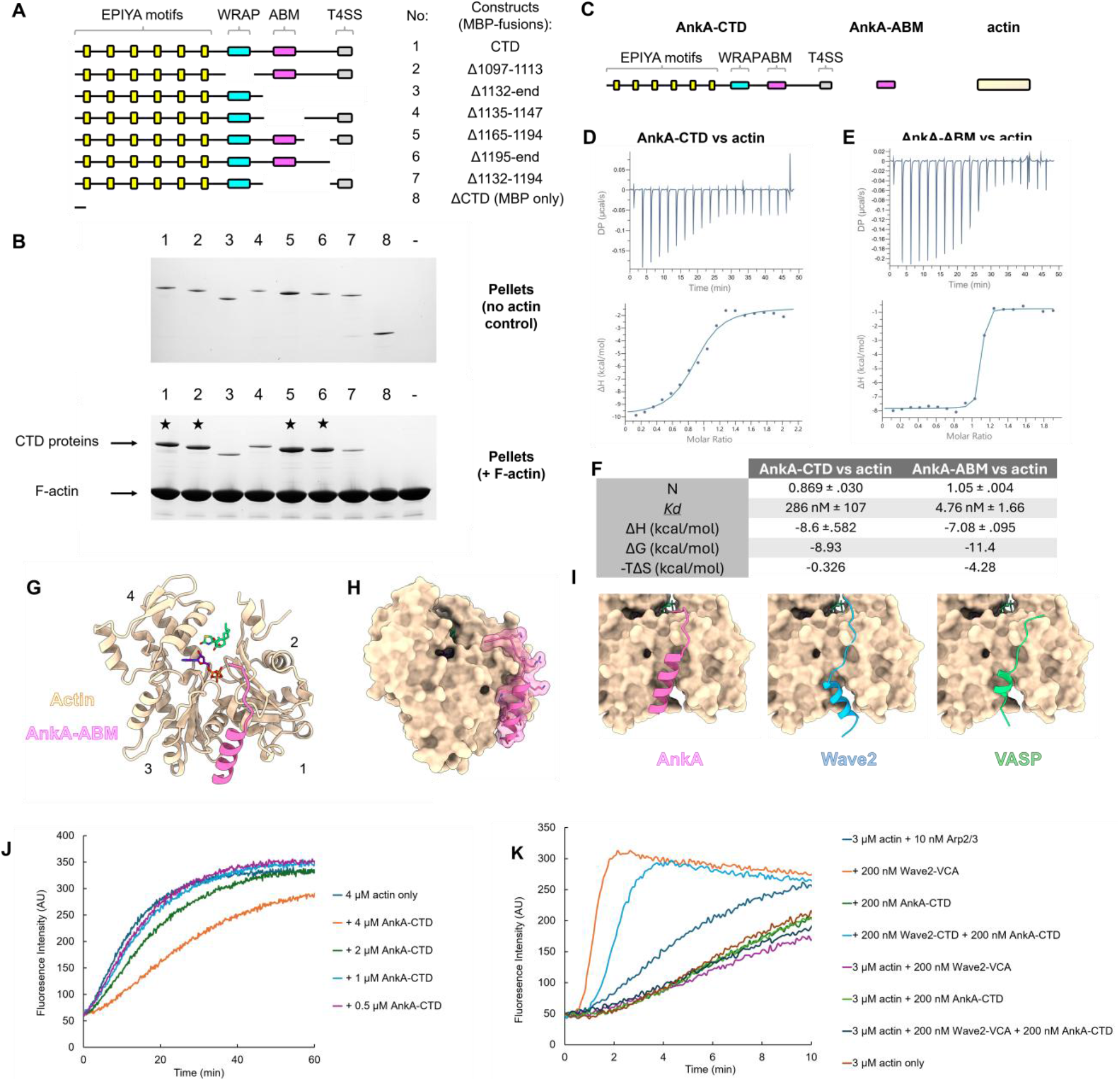
AnkA interacts with actin via a discrete Actin Binding Motif and inhibits actin polymerization *in vitro*. A. Schematic of the AnkA protein constructs used in F-actin pelleting assays. All were produced as MBP fusions. ABM: Actin Binding Motif. B. SDS-PAGE analysis of F-actin pelleting assays. Some AnkA-CTD proteins co-pelleted with F-actin (marked by a star) whereas others did not. C. Schematic of proteins and peptides used in the ITC analyses. Representative isotherms and integrated heats resulting from the titration of D. AnkA-CTD into G-actin, and E. a peptide based on the predicted actin binding motif of AnkA (AnkA-ABM) into G-actin. Results are representative of two independent experiments. F Dissociation constants and thermodynamic parameters extracted from ITC experiments. G. Cartoon representation of the AnkA-ABM:actin complex crystal structure solved to a resolution of 2.84 Å. The sub-domains of actin are numbered,1-4. AnkA-ABM binds within the cleft formed between sub-domains 1 and 3 at the barbed end of actin. H. Surface representation of the AnkA-ABM:actin complex demonstrating shape complementarity. I. Comparisons between AnkA-ABM:actin and the published structures of Wave2:actin (PDB:2A40; [41]) and VASP:actin (PDB: 2PBD, [68]). J. Actin-pyrene was used in actin polymerisation assays in which actin polymerisation was initiated by the addition of KCl, Mg^2+^, and ATP. The addition of AnkA-CTD protein caused a decrease in the rate of actin polymerisation. K. Actin pyrene assays including Arp2/3 and the Wave2-VCA protein. The addition of Arp2/3 plus Wave2-VCA initiated rapid actin polymerisation which was inhibited by the inclusion of AnkA-CTD protein. Results are representative of two independent experiments.

We next assessed whether AnkA-CTD could also bind to monomeric G-actin using ITC (**Fig. 4C**). Titration of AnkA-CTD (lacking any affinity tags) into G-actin yielded isotherms that were fitted to a one-binding site model, yielding a K_d_ of 286 nM, a stoichiometry close to 1:1, and binding being primarily enthalpy driven ((**Fig. 4D & F).** Following this, binding of a 35-amino acid peptide corresponding to the AnkA-ABM and flanking sequences (AnkA residues 1132-1166) to G-actin was assessed (**Fig. 4E**). The AnkA-ABM peptide bound to G-actin with a K_d_ of 4.76 nM and a stoichiometry also close to 1:1, indicative that this peptide constitutes the residues necessary for AnkA to bind actin with high affinity (**Fig. 4F).**

The binding affinities of actin binding peptides for actin are typically in the nM to low µM K_d_ range [41]. A 21 amino acid peptide based on the *Chlamydia* effector protein Tarp (which, like AnkA, also has EPIYA motifs) binds actin with a K_d_ of 102 nM [42] and a 31 amino acid WAVE2 peptide binds actin with a K_d_ of 52 nM [41]. Since the 34 amino acid AnkA-ABM peptide bound to actin with a higher affinity than WAVE2 it is possible that AnkA might compete with WAVE2 for binding to actin.

### Crystal structure of an AnkA-ABM peptide in complex with actin reveals molecular mimicry by AnkA

To determine the structural basis of AnkA interactions with actin we co-crystallized the Ank-ABM peptide with actin in the presence of ATP, Ca^2+^, and latrunculin B (which was included to inhibit actin polymerization during the time course of crystallization trials). The complex crystallized in space group *P* 4_3_ 2_1_ 2, x-ray diffraction data was collected to 2.84 Å, and the structure was solved by molecular replacement. The asymmetric unit contained three molecules of actin; each bound to latrunculin B, ATP, and Ca^2+^ (**Fig. 4G**, **Supplementary table 2**). The majority of actin could be modelled with residues at the N- and C-termini and residues corresponding to the D-loop (which is typically disordered in actin crystal structures) being absent. Helical density, not accounted for by actin, was present in the barbed-end clefts of all three actin protomers. For two of the three protomers in the ASU this density was poorly defined and lacked obvious protein side chains but was much more clearly defined in the third protomer (Chain A), likely due to crystal packing restricting the movement of the AnkA-ABM peptide. Accordingly, we were able to manually model 18 residues of the AnkA-ABM peptide (SSFAAELQAQRGKLRPVK), into the electron density map, yielding an AnkA-ABM:actin crystal structure (**Fig. 4G).**

In the complex, the AnkA-ABM peptide forms a 12-residue amphipathic alpha-helix that binds between actin subdomains 1 and 3. An additional 5 amino acids of peptide meander across sub-domain 1 towards the nucleotide binding site. Binding of AnkA-ABM is primarily mediated by shape complementarity (**Fig. 4H).** Residues F1136 and L1140 at the N-terminus of the AnkA-ABM helix extend into a hydrophobic pocket formed by actin residues Y143, I345, L346, and L349. Hydrogen bonds are formed between the sidechains of the AnkA helix residues E1139, Q1141, Q1143, and R1144, and actin residues T351, G146, S348, and A144, respectively. In the meander region, AnkA residue L1147 extends towards actin residue I345 and hydrogen bonds between the backbone of AnkA residues A1154 and P1155 and actin residues G23 and D25 stabilize the complex.

Comparisons between the AnkA-ABM:actin complex structure and those of WH2-domain peptides in complex with actin reveal a common binding mechanism (**Fig. 4I**) and a multiple alignment of selected WH2 motifs with AnkA-ABM reveals the presence of conserved or equivalent residues including hydrophobic helix residues and an LKKV-like sequence (**Supplementary fig. 1**). These results demonstrate that the AnkA-ABM is a WH2-like domain that binds to the barbed-end cleft of actin. Binding of AnkA to actin is expected to be mutually exclusive with the binding of other WH2 motifs, such as that of WAVE2.

### AnkA inhibits actin polymerization

Since AnkA binds to actin we next assessed whether this interaction has an effect on actin polymerisation. Accordingly, we measured the effects of AnkA on chemically induced actin polymerization using *in vitro* actin-pyrene polymerization assays (**Fig. 4J)**. The addition of KCl, Mg^2+^ and ATP to actin caused a rapid increase in actin polymerization. Titration of AnkA-CTD protein into identical assays resulted in a dose-dependent inhibition of actin polymerization. Some WH2 motifs are known to enhance actin polymerisation at low concentrations and to disrupt or inhibit at high concentrations [43]. We recorded no enhancement of actin polymerisation by AnkA-CTD at molar ratios down to 0.125:1. Use of the AnkA-ABM peptide in place of AnkA-CTD led to similar levels of actin polymerisation inhibition (**Supplementary fig.2**) We also assayed the impact of AnkA-CTD on pre-formed F-actin filaments but were unable to detect any effects. These results indicate that the AnkA can inhibit actin polymerization and that the AnkA-ABM alone is sufficient to elicit this effect, suggesting that AnkA sequesters actin monomers and possibly blocks actin barbed ends.

The WRC, via the WAVE2-VCA domain, binds to actin monomers and activates the Arp2/3 complex to initiate branched actin polymerisation. Since AnkA binds both the WRC and actin, we next questioned whether AnkA might inhibit actin polymerization downstream of WRC-activated Arp2/3 **(Fig. 4K)**. As expected, the inclusion of Arp2/3 plus WAVE2-VCA protein in actin-pyrene polymerization assays caused rapid actin polymerization compared to assays including Arp2/3 alone. The addition of equimolar concentrations (relative to WAVE2) of AnkA-CTD in these assays resulted in partial inhibition of WAVE2-VCA-activated Arp2/3-mediated actin polymerization. Collectively, these results demonstrate that AnkA inhibits actin polymerization *in vitro*, including when combined with WAVE2-activated Arp2/3, most probably by blocking of actin barbed-ends.

### AnkA and WRC components co-localise during infection

We sought to determine how AnkA protein localises during infection of human cells. Uninfected HL-60 cells and cells asynchronously infected with *A. phagocytophilum* were stained for confocal immunofluorescence microscopy using anti-AnkA antibodies. Whilst little to no AnkA staining was observed in uninfected control cells (**Supplementary fig. 3**), we observed varied localisation patterns of AnkA in *A. phagocytophilum* infected cells (**Fig. 5A)**. In some cells there was very little detectable AnkA signal despite the presence of an ApV (**Fig. 5A, top panels)**. In others, AnkA was present diffusely throughout the cytoplasm and nucleus (**Fig. 5A, middle panels**). In heavily infected cells, AnkA strongly co-localised with the ApV (**Fig. 5A, bottom panels)**.

**Figure 5.**
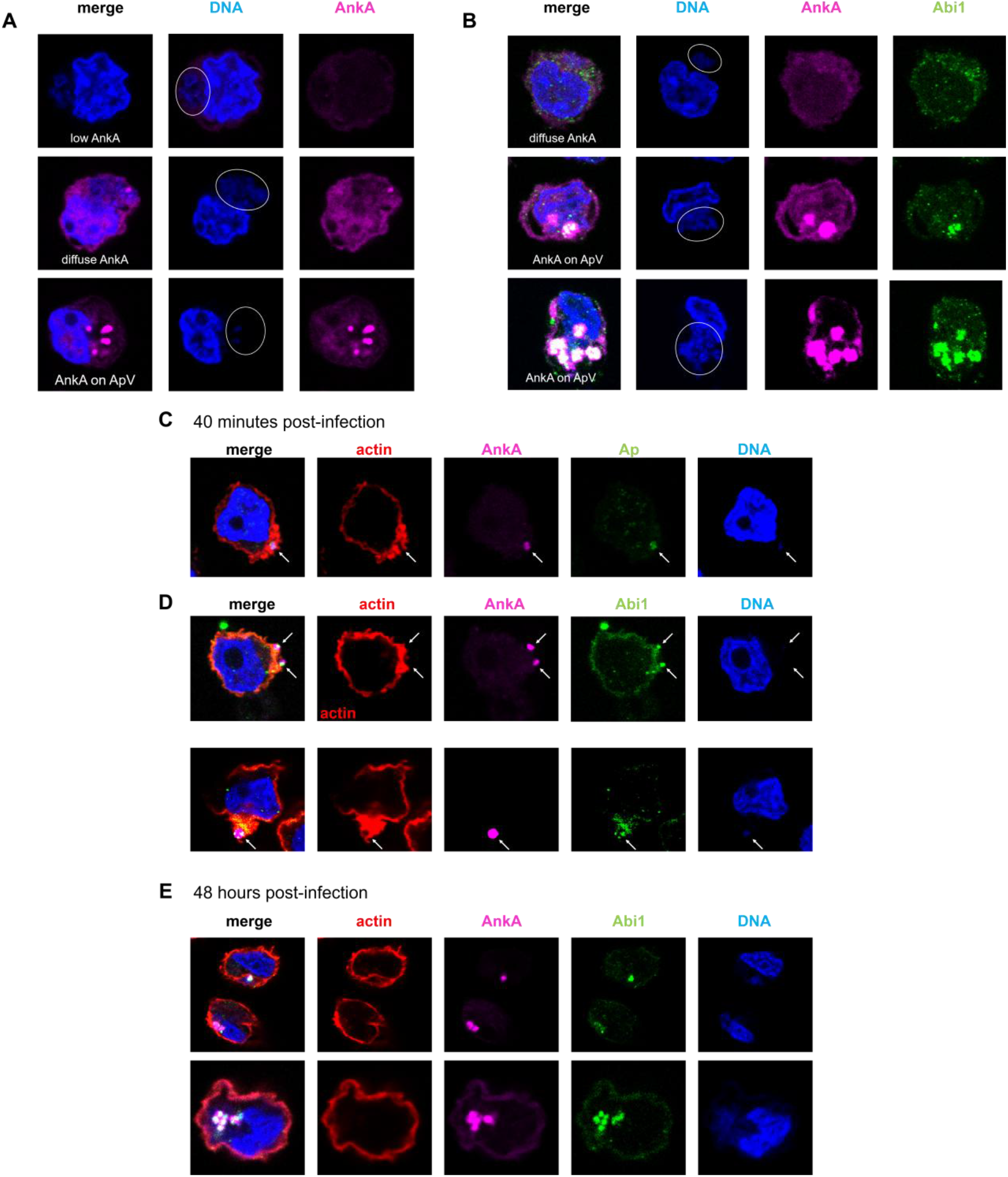
AnkA and Abi1 localisation during *A. phagocytophilum* infection of HL-60 cells. A-B: AnkA localisation and co-localisation with Abi1 in HL-60 cells asynchronously infected with *A. phagocytophilum*. A. HL-60 cells infected with *A. phagocytophilum* were fixed and stained for confocal immunofluorescence microscopy using anti-AnkA and Hoechst 33342. Host cells contained varied quantities of AnkA and different localisation patterns. B. Infected cells were also stained using anti-Abi1. AnkA and Abi1 co-localised prominently to large *A. phagocytophilum* inclusions. C-E: AnkA and Abi1 localise with *A. phagocytophilum* during the early and late stages of infection. HL-60 cells were infected with cell-free *A. phagocytophilum* for 40 minutes prior to fixing and staining for confocal immunofluorescence microscopy. C. *A. phagocyophilum* infected HL-60 cells stained using anti-*Ap* and anti-AnkA antibodies, phalloidin, and Hoechst 33342. D. Cells from the same infection stained with anti-Abi1 in place of anti-*Ap.* White arrows point to *A. phagocytophilum* bound to host cells. E. Infected cells at 48 hours post infection. White arrows point to *A. phagocytophilum* bound to host cells.

Next, we co-stained HL-60 cells with anti-AnkA and anti-Abi1 to determine whether Abi1 signal coincides with that of AnkA. Abi1 has non-specific cell-wide distribution in uninfected cells (**Supplementary fig. 3**) but co-localises with AnkA at the ApV in cells heavily infected with *A. phagocytophilum* (**Fig. 5B, middle and bottom panels**). Abi1 did not localise to the ApV in infected cells where AnkA was diffusely distributed in the cytoplasm, suggesting co-recruitment of AnkA and Abi1 (**Fig. 5B, top panel).** Staining of infected cells with anti-WAVE2 revealed a localisation pattern similar to that of Abi1 (**Supplementary fig. 4),** indicating that other components of the WRC are also recruited to the ApV with Abi1 and AnkA. The co-localisation of AnkA, Abi1, and WAVE2 late in infection suggest that AnkA is exposed to the host cytosol on the ApV and recruits the WRC.

Having identified that AnkA and WRC components localise to mature *A. phagocytophilum* inclusions, we next asked whether the two proteins associate during the earliest stages of infection, and whether this association correlates with perturbations of the actin cytoskeleton. Uninfected HL-60 cells were exposed to cell-free *A. phagocytophilum*, incubated for 40 minutes, then fixed and stained for immunofluorescence microscopy with anti-*Ap* antibody, anti-AnkA, and phalloidin (**Fig. 5C).** This is sufficient time for *A. phagocytophilum* to adhere to host cells and for some, but not all, to be internalised [44]. Single *A. phagocytophilum* cells bound to or entering HL-60 cells could be visualised in this manner with AnkA, visualised as a bright spot, coinciding with these bacteria surrounded by F-actin protrusions (**Fig. 5C)**. Staining cells for AnkA and Abi1 revealed that Abi1 also co-localises with *A. phagocytophilum* being engulfed and internalised (**Fig. 5D**).

Since the dense localisation of WRC at the ApV during late infection might be expected to coincide with a noticeable deposition of F-actin, we next visualised AnkA, Abi1 and actin in HL-60 cells at 48 hours post-infection, when numerous large inclusions would be present (**Fig. 5E)**. In line with our biochemical data demonstrating that AnkA can inhibit actin polymerization, in most cases we did not see F-actin rich structures coinciding with AnkA and Abi1 at the ApV. This is also in line with previous studies that demonstrated a lack of actin structures around the ApV in RF/6A cells (a rhesus monkey endothelial cell line) and uncertainty regarding whether F-actin is specifically assembled around the ApV [44]. One caveat is that HL-60 cells are small and rounded, making fine structures difficult to resolve by conventional confocal microscopy, and fixed cells provide a snapshot. Finer and transient F-actin structures may therefore have escaped detection.

In total, immunofluorescence data indicate that AnkA, the WRC, and F-actin co-localise early during infection during bacterial attachment and entry. AnkA distribution in cells with established infection is diverse but, during the later stages of infection, AnkA and the WRC robustly localise to the inclusion in the absence of F-actin judged by phalloidin staining. Whilst our data do not resolve whether AnkA staining at the inclusion is of secreted protein into the host cell or AnkA within bacterial cells, the co-localisation of its binding partner, the WRC, at the inclusion is strongly indicative that at least some AnkA had been secreted and was exposed to the host cytosol. The apparent conflict between our *in vitro* actin polymerisation data demonstrating AnkA inhibits actin polymerisation and the co-localisation of AnkA and the WRC with F-actin during early infection are discussed below (see discussion).

### Candidate WH2 and WRAP-binding motifs in the effector proteins of other obligate intracellular bacteria

We next asked whether AnkA-WRAP and AnkA-ABM motifs occur in the effectors of other bacteria. Genus *Anaplasma* includes species tropic for distinct host blood cells such as erythrocytes, platelets, and granulocytes. WRAP, ABM, and EPIYA motifs and their relative positions in the AnkA-CTD are generally conserved in the AnkA proteins of *A. phagocytophilum*, including isolates or strains originating from the USA, Europe and Asia and from diverse hosts (**Fig. 6A, Supplementary fig. 5A)**. Of those analysed, only one *A. phagocytophium* isolate (AKS2020 from the Himalayan marmot) encodes an AnkA that lacks candidate WRAP and ABM motifs. Two tandem candidate ABM and WRAP motifs, and multiple EPIYA motifs, were identified in the Ank-CTD of *A. platys*, a species which infects platelets (**Fig. 6A, Supplementary fig. 5B)**. The AnkA of *A. bovis,* which infects monocytes, has EPIYA-like and ABM-like motifs but lacks a WRAP motif. In contrast, WRAP, ABM, and EPIYA motifs are absent from the AnkA proteins of erythrocyte-infecting *Anaplasma* species such as *A. marginale, A. ovis, and A. capra.* This suggests that WRC binding, actin binding, and EPIYA-mediated SH2 binding are functionally linked attributes of *Anaplasma* AnkA proteins: WRAP motifs are found alongside ABM- and EPIYA-motifs or not at all. These attributes correlate with *Anaplasma* species that infect neutrophils or platelets, cells that can form lamellipodia, produce secretory granules, and be regulated or activated by non-receptor tyrosine kinases, but not with species that infect erythrocytes, which lack these features.

**Figure 6.**
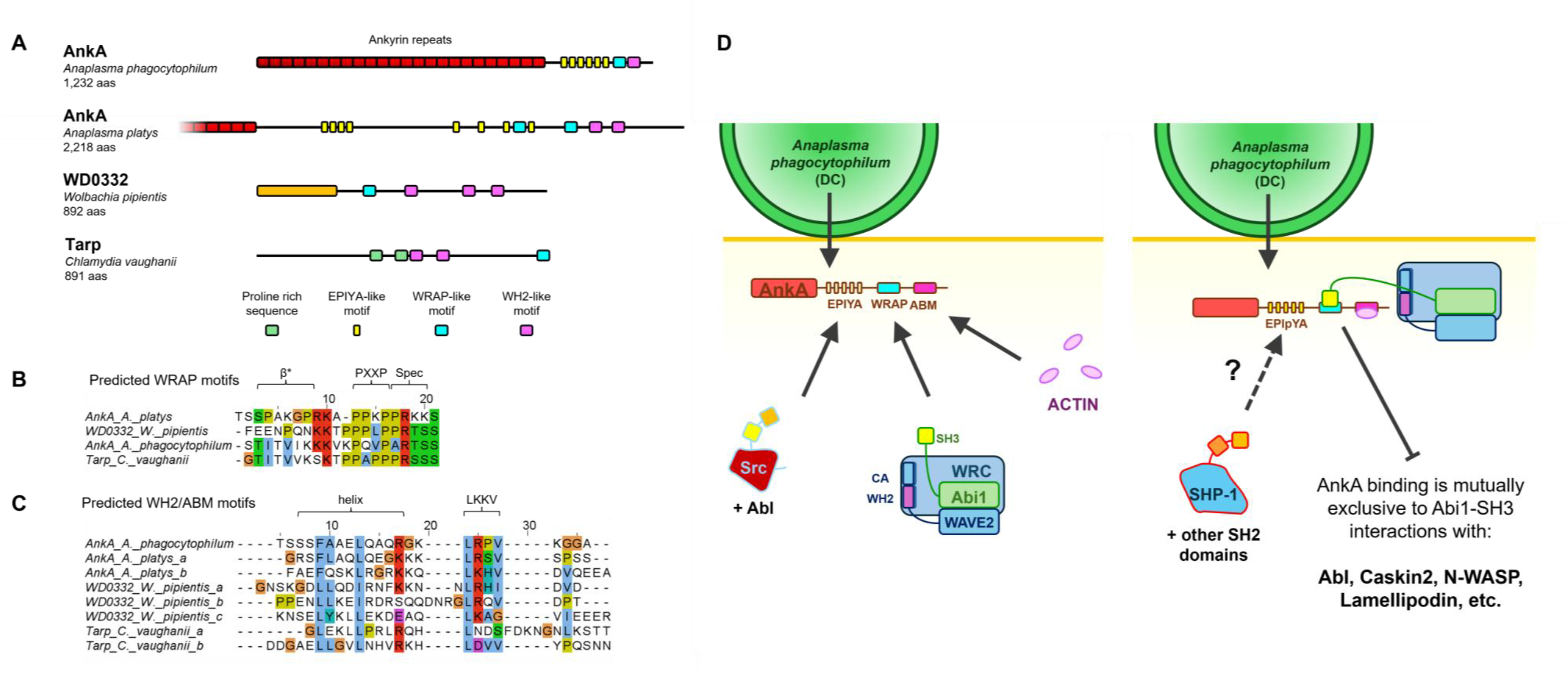
Identification of AnkA-WRAP-, ABM-, and EPIYA-like motifs in probable effector proteins and a model for AnkA functions. A. Schematic of protein domains. Protein regions that are mostly predicted to be disordered are shown as a solid black line. The ankyrin repeat domain of *A. platys* AnkA (shown as red blocks and of similar size to that of *A. phagocytophilum*) is only partially shown due to the large total size of the protein. An uncharacterized domain of WD0332 is shown as an orange block. B. Sequences of AnkA-WRC-like motifs. C. Sequences of WH2-like motifs. Conserved residues are coloured according to residue type according to Clustal Omega colouring scheme. D. Cartoon summary of AnkA functions during early stages of infection.

We extended our search to other bacteria, focussing on known or predicted effector proteins. WRAP-like motifs were found in a hypothetical protein conserved in *Wolbachia,* an obligate intracellular bacterium of the *Anaplasmataceae* family which infects an estimated 40% of terrestrial arthropods [45]. This protein, called WD0332 in *Wolbachia pipientis* wMel, is a predicted secreted effector protein with WH2-like motifs [46]. Both candidate WH2- and WRAP-motifs are conserved in this predicted effector protein across many *Wolbachia* genomes (**Fig.6A & B)**. A WRAP-like motif was also found in the effector Tarp from the obligate intracellular bacterium *Chlamydia vaughanii* and then (by using this motif as a search term) in the Tarp proteins of *Chlamydia ibidis* and *Chlamdia felis*. (**Fig. 6A & B)**. Whilst the Tarp protein of *Chlamydia trachomatis* has actin binding- and EPIYA-motifs and has a defined role in remodelling host actin during infection [47, 48] the Tarp proteins of most other *Chlamydia* lack EPIYA-motifs but retain actin binding motifs and have not been studied in detail.

The analyses suggest that the ability of *Anaplasma* to subvert actin and the WRC via AnkA correlates with host cell tropisms, being generally present in *Anaplasma* species which infect host cells that are reliant upon WRC-driven processes for their normal functions in mature cells. The co-incident presence of WH2 and AnkA-WRAP-like motifs in the disordered regions of predicted or proven effector proteins from other obligate intracellular bacteria suggest that AnkA might represent a larger family of bacterial effector proteins.

## Discussion

In this study we have discovered a new role for the AnkA effector of *A. phagocytophilum* in direct and indirect targeting of the actin cytoskeleton. Multiple pathogens are known to target the Rac-WRC-Arp2/3 signalling axis to facilitate infection and bacterial effector proteins have been identified that target Rac and Arf GTPases upstream of the WRC [31, 49], the downstream Arp2/3 complex [50–52]; and other WASP-family proteins that function in parallel with the WRC [53, 54]. The AnkA of *A. phagocytophilum* is, to our knowledge, the first example of a bacterium directly targeting the WRC and therefore represents a newly identified strategy for bacterial subversion of the host cytoskeleton.

Based on our structural data, the binding of AnkA to the Abi1-SH3 component of the WRC is mutually exclusive with the ability of the latter to bind some of its endogenous binding partners. Thus, we expect that AnkA would block the interaction of Abi1-SH3 with Abl and other proteins known to bind Abi1-SH3 such as Caskin2, N-WASP, and Lamellipodin [40, 55–57]. In this way, AnkA likely influences WRC activity by selectively competing with a specific subset of WRC interaction partners. Intriguingly, AnkA binding to Abi1 is reasoned to facilitate the formation of a tripartite AnkA:Abi1:Abl complex [13]. However, since Abi1 binding to Abl kinase is dependent on the latter engaging the Abi1-SH3 domain [39], for such a complex to exist it must form independently of the normal Abi1-SH3: Abl interface.

Via a WH2-like motif, AnkA binds actin and inhibits its polymerization, including downstream of the Arp2/3 complex, *in vitro*. This function of AnkA might allow localised dampening of actin polymerisation during infection, such as at the ApV where AnkA and Abi1 accumulate but in the apparent absence of F-actin structures. In accumulating at the ApV, AnkA might influence actin-dependent vesicular traffic and sequester cellular WRC.

Since it is currently not possible to engineer point mutations in *A. phagocytophilum* and AnkA is likely to be essential for this bacterium [13, 20], dissecting precisely how this effector supports intracellular survival and contributes to pathogenesis is challenging. Nonetheless, our findings suggest three phases of AnkA function in the vertebrate host (**Fig. 6D, Supplementary fig. 6)**. First, AnkA engagement of the WRC during early infection - coinciding with F-actin protrusions - may support ApV biogenesis, given the WRC role in endocytic pathways. Second, during bacterial proliferation, AnkA distributes throughout the host cell, where it influences the nucleus and other signalling pathways, as described elsewhere. Finally, in later infection, dense clustering of AnkA and the WRC at the ApV suggests that AnkA sequesters cellular WRC and dampens actin polymerisation at the ApV prior to bacterial release. A caveat of our model is that AnkA inhibits actin polymerization *in vitro*, including downstream of the Arp2/3 complex, yet F-actin protrusions co-localise with AnkA and the WRC during entry. Our *in vitro* assays tested AnkA activity against actin and Arp2/3 in the absence of the intact WRC, its upstream activators, or other influencing factors, which would be present *in vivo*. This may account for the apparent tension between our observations *in vitro* and during bacterial entry. Further work is required to discern how location, concentration, and post-translational modifications govern AnkA activity at distinct phases of infection.

Taken together, the results of this study reveal substantive mechanistic insights into multi-factorial actin subversion by *A. phagocytophilum* and a newly identified strategy for targeting the host actin cytoskeleton by a microbe in direct targeting the WRC and actin. Since the WRC has a central role in many cellular processes that are central to phagocyte functions such as cell migration, phagocytosis, and the assembly of phagocyte oxidase [58], and since these processes are disrupted by *A. phagocytophilum* infection, it is possible that AnkA has wide-ranging impacts on host cell biology. Furthermore, that AnkA and WRC co-localise during bacterial internalization and during the late stages of infection suggest that AnkA plays a role in *A. phagocytophilum* colonization of host cells and dissemination. The presence of WRAP and ABM motifs in other obligate intracellular bacteria, as well as recent work by others demonstrating that the apicomplexan pathogen *Toxoplasma* targets the WRC via a different mechanism [59, 60], raises the possibility that multiple other pathogens have independently evolved similar strategies for targeting WRC.

## Methods

### Bacterial and eukaryotic cell culture

*Escherichia coli* strain DH5α was used for routine molecular biology and for plasmid preparations whereas *E. coli* strain BL21(DE3) was used for overexpression of recombinant proteins for purification. Human cell line HEK293 was cultivated in DMEM medium supplemented with 10% v/v fetal bovine serum (FBS) and 2 mM Glutamax.

The human HL-60 cell line, used for routine cultivation of *A. phagocytophilum* Webster and for infection experiments, was cultivated in RPMI 1640 medium supplemented with 10% v/v FBS, 2 mM Glutamax, and 1 mM Pyruvate. *A. phagocytophilum* was routinely cultivated as asynchronous infections of HL-60 cells as described elsewhere [61].

### Protein expression and purification of AnkA protein for polyclonal antibody generation

DNA sequences encoding AnkA (APH_0740) residues 1-588 were amplified from *A. phagocytophilum* HZ gDNA by PCR and cloned into a modified pET41c vector (lacking the glutathione S-transferase sequences) by a restriction-free cloning approach as described elsewhere [62, 63] This yielded an expression vector encoding AnkA residues 1-588 (corresponding to over half of the ARD of AnkA) fused to a C-terminal His_8_ tag. For purification of the resulting recombinant protein (which we term AnkAT2) *E. coli* cells were grown in Terrific Broth supplemented with kanamycin at 37 °C until an optical density at 600 nm of ∼0.6 had been reached. Gene expression was induced by the addition of 0.5 mM IPTG for 20 hours at 20 °C. Harvested cells from 2L of culture were resuspended in Buffer A (50 mM Tris-HCl, pH 7.5; 20 mM imidazole-HCl, pH 7.5; 400 mM NaCl; 2 mM beta-mercaptoethanol) and then sonicated. Cell lysates were clarified by centrifugation at 80,000 x g for 45 minutes and the resulting lysate was applied to a 5 ml Hi-Trap His Column (Cytiva). Non-specifically bound proteins were washed from the column with 20 column volumes of Buffer A prior to stepwise elution of AnkAT2 with Buffer A supplemented with 5, 10, 20 and 100 % Buffer B (as Buffer A but with 400 mM imidazole-HCl, pH 7.5). Fractions containing AnkAT2 (typically those eluting at 20% Buffer B) were pooled and dialysed against Buffer C (20 mM Bicine-NaOH, pH 8.5; 250 mM NaCl; 1 mM DTT) prior to gel filtration using a Superdex 200 26/600 column (Cytiva) in Buffer C. Peaks containing pure AnkAT2 protein were pooled, concentrated by ultra-filtration to ∼ 10 mg/ml, and then transferred into PBS on a HiPrep 26/10 Desalting column immediately prior to flash freezing in liquid nitrogen. AnkAT2 protein prepared in this way was used for polyclonal antibody generation and purification.

### Polyclonal antibody generation and purification

AnkAT2 protein in PBS was used for polyclonal antibody generation in rabbits as a service provided by Eurogentec. To purify AnkAT2 antibodies, AnkAT2 protein was coupled to CNBr Activated Sepharose 4B (Cytiva) according to the manufacturer instructions to yield an AnkAT2 affinity column. Antibodies were purified from serum using AnkAT2 protein covalently coupled to CNBr-activated Sepharose 4b.

### Immunoprecipitation of AnkA from infected cell lysates

For immunoprecipitation of AnkA from infected cells, anti-AnkA antibodies were coupled to Dynabeads using the Dynabeads Antibody Coupling Kit (Invitrogen), following manufacturer instructions. Uninfected and *A. phagocytophilum* infected HL-60 cell cultures were used to prepare lysates for immunoprecipitation. Cells were harvested by centrifugation (200 rpm for 5 mins), washed with PBS, then frozen at −80 °C before use. An equal number of infected and uninfected cells (approximately 1.2 x 10^8^ cells for each) were lysed in IP buffer (20 mM Tris-HCl, pH 8.0; 100 mM NaCl; 1 mM DTT; 1 % v/v nonidet P-40; 2 mM EDTA; Complete EDTA-free Mini Protease Inhibitors) by sonication. Lysates were clarified by centrifugation at 13,000 rpm for 10 minutes and the resulting supernatants were incubated with anti-AnkA dynabeads for 90 minutes at 4 °C. Beads were separated from lysate using a magnetic rack and then sequentially washed with IP buffer containing 0.1 % v/v nonidet P-40. Beads were washed once with IP buffer lacking any detergent and then transferred to a clean microfuge tube. To minimise the presence of antibody fragments in the resulting IP samples, bound proteins were eluted with sequential washes with 0.1 N ammonium hydroxide plus 0.1 mM EDTA. Eluted proteins were dried by vacuum centrifugation, resuspended in reducing SDS-PAGE gel loading dye, then run ∼ 1 cm into SDS-PAGE gels prior to staining with Coomassie. Lanes were excised (typically 0.5 cm x 0.5 cm) were then sent for LC-MS-MS analysis at St Andrew’s University (UK).

### Immunoprecipitations of GFP and GFP-tagged proteins from HEK293 cells

The AnkA ORF (*A. phagocytophilum* strain HZ, APH_0740) was cloned into the vector pcDNA3.1-C-GFP by restriction-free cloning. The resulting plasmid, encoding the AnkA protein with GFP fused at the C-terminus of the protein, was used for Q5 mutagenesis (New England Biolabs) to yield an additional vector encoding the AnkA-ARD (residues 1-859) fused to eGFP. These vectors and the empty parental vector encoding eGFP only were used for GFP-Trap IP experiments. Briefly, expression vectors were transfected into HEK293 cells using FuGene HD Transfection Reagent according to the manufacturer’s instructions (Promega) which were then harvested at 48 hours post-transfection. Cells were washed with PBS and then used for immunoprecipitations with GFP-Trap agarose beads (Chromotek) according to the manufacturer instructions. Proteins were released from the beads by boiling in SDS-PAGE loading dye, then run a short distance into SDS-PAGE gels prior to excision and sending for LC-MS-MS analysis.

### Protein expression and purification of GFP-tagged and untagged AnkA-CTD proteins

DNA sequences encoding AnkA residues 890-1232 were amplified by PCR and cloned in a similar manner to that described above. The destination plasmid was a modified pET41c vector, lacking the GST-sequences but encoding a tobacco-etch virus (TEV) protease cleavage site, eGFP, and an His_8_ tag. This yielded an over-expression vector encoding the AnkA-CTD fused to a TEV protease site, eGFP, and His_8_ at the C-terminus. Overexpression and nickel purification of the recombinant protein was as described for AnkAT2 but with the addition of Complete EDTA-free protease inhibitors (Roche) during cell lysis. Peaks of the AnkA-CTD-TEV-eGFP-His_8_ protein that eluted from the HiTrap His column were dialysed into Buffer D (20 mM Tris-HCl, pH 8.0; 50 mM NaCl, 2 mM beta-mercaptoethanol) and then further purified by anion exchange with CAPTO Q columns (Cytiva), using Buffer D and a gradient of 50 mM to 1 M NaCl. This yielded pure AnkA-CTD-TEV-eGFP-His_8_ protein which was used in pull-down experiments.

To generate untagged AnkA-CTD protein, a portion of AnkA-TEV-CTD-eGFP-His_8_ was treated with TEV protease at 4 °C overnight to cleave the C-terminal fusion proteins from AnkA-CTD. The TEV cut protein preparation was passed through a HiTrap His column to remove TEV-eGFP-His_8_. The flow-through from this column (containing untagged AnkA-CTD protein) was concentrated and then further purified by gel filtration using a Superdex 200 26/600 column equilibrated in Buffer D supplemented with 200 mM NaCl. Peaks containing untagged purified AnkA-CTD were pooled and concentrated to ∼15 mg/ml. Untagged recombinant AnkA-CTD prepared in this way was used for ITC experiments.

### Protein expression and purification of His_6_- and MBP-tagged Abi1 proteins

The Abi1 coding sequence was cloned by restriction-free cloning into a modified pET28a vector encoding N-terminal His_6_, MBP, and TEV protease sequences. This yielded an expression construct encoding His_6_-MBP-TEV-Abi1 protein. Q5 mutagenesis was used to create deletion mutants of His_6_-MBP-TEV-Abi1 corresponding to Abi1 residues 1-157 (Abi1-NTD), 317-489 (Abi1-CTD), and 404-489 (Abi1-SH3) domains. These resulting vectors were used for protein over expression and purification using HisTrap columns as described above with proteins being dialysed into Buffer D prior to gel filtration on Superdex 200 26/600 columns.

### Pull-down assays

Pull-down assays using His_6_-MBP-tagged Abi1 proteins as bait were achieved by incubating MBP-tagged proteins with Amylose Resin (New England Biolabs) equilibrated in Buffer D supplemented with 200 mM NaCl. Unbound proteins were removed by washing the resin with the same buffer and then the beads were incubated with a 2-fold molar excess of AnkA-GFP protein for 1 hour at room temperature with gentle mixing. The beads were washed three times, then collected by low-speed centrifugation prior to boiling in SDS-PAGE loading dye. Samples were analysed by SDS-PAGE.

### Overexpression and purification of untagged Abi1-SH3 protein

The Abi1-SH3 domain was expressed and then purified on HisTrap columns as described above. Following this purification step, the protein was supplemented with TEV protease and dialysed against Buffer D overnight to cleave the fusion protein. Cleaved protein was passed through a HisTrap column to remove the His_6_-MBP tag and the flow-through, containing untagged Abi1-SH3, was concentrated and further purified by gel filtration with a Superdex 75 26/600 column in Buffer D supplemented with 200 mM NaCl. The resulting protein was concentrated to ∼16 mg/ml and used for ITC and crystallography experiments.

### Isothermal calorimetry

ITC experiments were run in a PEAQ-ITC Instrument (Malvern). Peptides were synthesized by Peptide2.0 with N-terminal acetylation and C-terminal amidation modifications to better mimic the native proteins they were designed from. Proteins were extensively dialysed against a common buffer prior to use in ITC experiments whereas lyophilized peptides were resuspended in this same buffer.

For experiments using untagged Abi1-SH3, untagged AnkA-CTD, and the AnkA-WRAP peptide, the buffer used was 20 mM Tris-HCl, pH 8.0; 200 mM NaCl. The AnkA-WRAP peptide (500 µM) was titrated into Abi1-SH3 (55 µM). The Abi1-SH3 protein (400 µM) was titrated into AnkA-CTD (25 µM). The latter experiment was performed in this way as it was challenging to produce sufficient untagged AnkA-CTD to achieve the high concentrations required for it to serve at the titrant.

Experiments using actin, untagged AnkA-CTD, and the AnkA-ABM peptide were performed in modified G-buffer (2 mM Tris-HCl pH 8, 200 μM ATP, 0.1 mM CaCl_2_). The AnkA-ABM peptide (130 µM) was titrated into actin (13 µM), and AnkA-CTD (90 µM) was titrated into actin (8 µM).

### Protein crystallography

Abi1-SH3 was co-crystallized with the AnkA-WRAP peptide by mixing Abi1-SH3 protein at 10 mg/ml with AnkA-WRAP peptide at a 1:1.2 molar ratio and then screening against Molecular Dimensions crystallography screens. Crystals were obtained in a MIDAS screen condition comprised of 30 % v/v glycerol ethoxylate and 0.1 M Tris-HCl pH 8.0. The Abi1-SH3:AnkA-WRAP complex crystallized in space group *C* 1 2 1. X-ray diffraction data was collected at Diamond Light Source Beamline I24 at 100K and at a wavelength of 0.999 Å. Data were processed with the xia2 3dii pipeline. Data were collected to 1.38 Å but, due to low completeness at high-resolution, were truncated to 1.69 Å and a new free R test set to yield completeness of greater than 80% in the high-resolution shell and an unbiased test set.

Actin was co-crystallized with the AnkA-ABM peptide in the presence of latrunculin B (to prevent F-actin formation), Ca^2+^, and ATP. Actin (7 mg/ml) in G-buffer and supplemented with 1 mM Latrunculin B, was mixed in a 1:2 molar ratio with AnkA-ABM peptide and incubated on ice for 30 minutes before being used in vapour diffusion experiments using Molecular Dimensions crystallography screens. The actin:AnkA-ABM complex crystallized in space group P 43 21 2 in the JCSG screen condition comprised of 0.2 M ammonium sulfate, 0.1 M BIS-Tris pH 5.5, 25 % w/v PEG 3350. X-ray diffraction data was collected at Diamond Light Source Beamline I24 at 100K, at a wavelength of 0.900 Å, and were processed with the xia2 3dii pipeline. Data were initially processed as *P* 4_1_ 2_1_ 2 but were reprocessed as *P* 4_3_ 2_1_ 2. Diffraction images showed ice rings at ∼ 3.0, ∼3.3, ∼3.8 Å but plots of completeness and CC1/2 were relatively unaffected in these regions (>99.5 and >0.4, respectively). Data were collected to a resolution of 2.84 Å. The high-resolution shell of the data had a low I/sigI value (0.5) but a CC1/2 value well above accepted cut-off thresholds (0.368), indicating the presence of statistically significant signal in this shell. As such, we trialled paired refinements with the full dataset and with truncated datasets. Using this approach, we found that excluding these high-resolution reflections did not lead to any improvement in model geometry, or R_free_, or in map quality. As such, the full dataset was used for solving the structure.

Diffraction data processing and refinement statistics are provided in Supplementary tables 1 and 2. The crystal structures of the Abi1-SH3: AnkA-WRAP and the actin:AnkA-ABM complex were solved by molecular replacement using a Colabfold model [64] of the Abi1-SH3 and the structure of Latrunculin B Bound to actin (extracted from PDB: 2Q0U; [65]) as respective search models and using Phaser in the Phenix software suite. In both cases, regions of the AnkA-WRAP and AnkA-ABM peptides were manually modelled in Coot. Structures were refined iteratively using Phenix Refine software and manual building in Coot. The PDB-Redo server was also used for refinement in the case of AnkA-ABM:actin [66].

### Transfection and confocal microscopy for lamellipodia and actin observations

B16F1 cells were grown in DMEM containing 10% fetal bovine serum and 4.0 mM of L-glutamine in 35 mm tissue culture Petri-dishes at 37 °C and 5%CO_2_. Approximately 70% confluent cells were transfected with 2.5 mg pEGFP-C1 or AnkA-eGFP, and pmCherry-C1 or pAbi1-mCherry expression plasmid vectors in separate experiments. After 24 hours of transfection, cells were trypsinised and plated on laminin coated glass bottom tissue culture dishes and allowed to adhere for 5 hours in CO_2_ incubator at 37 °C prior to live cell confocal imaging by Zeiss 880 AiryScan confocal microscope using 63x/1.4NA objectives. Cells were imaged in 37 °C and 5%CO_2_ enclosed box at an interval of 10 secs.

### Protein expression and purification of AnkA-MBP proteins

The full-length AnkA ORF, codon optimised for expression in *E. coli* was synthesized and cloned into the pMAL-c4X expression vector by Genscript to yield a construct encoding the full-length AnkA protein with MBP fused at the N-terminus. This plasmid was used as the template for Q5 mutagenesis (New England Biolabs) to delete AnkA residues 1-959, yielding a new construct encoding MBP-AnkA-CTD residues. This plasmid was then used as the template for further Q5 mutagenesis to yield constructs encoding AnkA-CTD proteins with deletions as detailed in Fig. 4A. Over-expression and lysis of *E. coli* expressing MBP-AnkA-CTD proteins was as described for AnkAT2 but with Ampicillin as the antibiotic selection. Initial purification of these proteins was achieved using MBP-Trap columns (Cytiva) following the manufacturer’s instructions. Following this purification step, MBP-AnkA-CTD proteins were purified by size exclusion chromatography as described for untagged AnkA-CTD protein. Proteins prepared in this way were used for pull-down and actin pelleting experiments.

### Actin purification, F-actin pelleting assays, and actin polymerization assays

Actin was purified from rabbit muscle acetone powder (Sigma-Aldrich) as described [67]. Pyrene-labelled actin and pure Arp2/3 complex were purchased (Cytoskeleton). Actin polymerization assays were performed according to protocols here [67] at concentrations marked in figures and figure legends. Actin polymerization assays using Arp2/3 complex were performed with proteins prepared and incubated in buffer (20 mM Tris-HCl, 50 mM KCl, 0.1 mM ATP, 1 mM MgCl_2_ buffer) for 15 minutes at 20 °C prior to the addition of actin stock. F-actin pelleting assays were performed by the method described in the Actin Binding Protein Spin-Down Assay Biochem Kit: rabbit skeletal muscle actin manual (Cytoskeleton). Wave2 for use in actin assays was expressed as an MBP-fusion protein and purified as described for Abi1 proteins.

### Immunofluorescence microscopy

Uninfected and *A. phagocytophilum* infected HL-60 cells were collected by centrifugation, washed in PBS, then resuspended in PBS at a density of 1.5-2.5 x 10^7^ cells per ml. These were then dispensed onto coverslips that had been pre-treated with 0.1% poly-l-lysine. After 10 minutes incubation on the slides, 4% PFA in PBS was added to the cells. After an additional 10 minutes incubation, the cells were washed three times with PBS to remove PFA and any cells that had not adhered to the coverslips. Cells were permeabilized by a 10-minute incubation with 0.1% Triton X-100 for 10 minutes and then blocked in 2% BSA in PBS at 4 °C. Fixed, permeabilized, cells were then labelled for immunofluorescence microscopy using rabbit anti-AnkA, mouse anti-Abi1 (Santa-Cruz Biotechnology), mouse anti-WAVE2 (Santa-Cruz Biotechnology), anti-HE (a gift from Prof Dumler), and fluorescently-labelled secondary antibodies (Invitrogen). Actin was stained with Alexa Fluor 555 Phalloidin and DNA was stained with Hoechst 33342 prior to mounting slides with ProLong Diamond Antifade mountant. For synchronous infections, cell-free *A. phagocytophilum* was produced following the protocol here [61].

Images were acquired with a Leica SP8 confocal system attached to a Leica DM I8 inverted microscope using an oil-immersion objective lens (63X, N.A. 1.4) and excitation at 405nm, 488nm, 561nm and 633nm. Imaging parameters were selected to optimise XY resolution and image stacks acquired at 0.5 mm intervals followed by maximum projection using Leica LASX software.

## Supporting information

Supplementary xls files

## Acknowledgements

We thank Drs Richard Logan (University of Birmingham), David Hardy (Aston University), Harriet Giddings (formerly of University of Birmingham), Jack Bryant (University of Nottingham), and Manuel Banzhaf (Newcastle University) for assistance during the early stages of this work. We would also like to thank Profs Kelly Brayton (Washington State University) and Robert Insall (University College London) for discussions.

This work was supported by funding from UKRI Future Leaders Fellowship MR/Y01748X/1, Academy of Medical Sciences Springboard Award SBF009/1204, and Royal Society Research Grant RGS\R1\231117. Preliminary data that led to these awards, essentially seeding this work in the conception stages, was generated with seed corn awards from the Cabot Institute for the Environment (University of Bristol), a Wellcome Trust Institutional Strategic Support Fund Global Mobility Award (managed by the University of Birmingham) and a Wellcome Trust Institutional Strategic Support Fund Award (managed by the Elizabeth Blackwell Institute, University of Bristol). Hannah Burge was funded by University of Bristol Faculty for Health and Life Sciences PhD scholarship. Joshua Lee was funded by a University of Bristol scholarship. Mass spectrometry carried out by the University of St Andrews Biomedical Sciences Research Complex Mass Spectrometry & Proteomics Core Facility was supported by BBSRC equipment grant BB/T017686/1.

The authors acknowledge the use of QED Science AI tool for pre-submission review and logic checks (https://www.qedscience.com). Claude AI was used for proof-reading (https://claude.ai). The authors take full responsibility for the final content and integrity of this work.

## Author contributions

Conceptualization: H.B, S.P.S., P.J.M., A.L.L., L.M., M.J., J.S.D., and I.T.C. Formal analysis: H.B., S.P.S., J.W., S.A.S., A.L.L., L.M., M.J., J.S.D., and I.T.C. Funding acquisition: I.T.C. Investigation: H.B, S.P.S., J.W., S.P., K.W., R.B., J.W.L., S.A.S., S.L.S., M.J., and I.T.C. Methodology: all authors. Resources: S.A.S., S.L.S., A.L.L., L.M., M.J., J.S.D., and I.T.C. Writing – original draft: H.B., S.P.S., and I.T.C. Writing – review & editing: H.B, S.P.S., K.W., J.W.L., P.J.M., A.L.L., L.M., M.J., J.S.D., and I.T.C.

## Data availability

Coordinates and structure factors for the AnkA-WRAP:Abi-SH3 and AnkA-ABM:actin structures generated in this study have been deposited in the PDB database under the accession codes 33SA and 33ZK, respectively.

## Ethics declarations

The authors declare no competing interests.

