## Supplementary figures and tables for "Direct targeting of the WAVE-regulatory complex and actin cytoskeleton by the *Anaplasma phagocytophilum* effector, AnkA"

**Supplementary files**

**Supplementary xls 1**

Results of LC-MS-MS analysis of AnkA immunoprecipitates from *A. phagocytophilum* infected and uninfected HL-60 cell lysates

**Supplementary xls 2**

Results of LC-MS-MS analysis of GFP immunoprecipitates from cell lysates of HEK293 cells expressing AnkA-3GFP and 3GFP

**Supplementary xls 3**

Results of LC-MS-MS analysis of GFP immunoprecipitates from cell lysates of HEK293 cells expressing AnkA-3GFP and AnkAΔCTD-3GFP

**Supplementary video 1:** Recruitment of AnkA-eGFP at the lamellipodia edge. B16F1 cells expressing AnkA-eGFP were allowed to adhere and migrate on laminin A coated glass bottom dishes and observed by AiryScan confocal microscopy. Filmed at 1 frame/10 seconds, movie shows 7frames/second.

**Supplementary video 2:** Recruitment of Abi1-mCherry at the lamellipodia edge. B16F1 cells expressing Abi1-mCherry were allowed to adhere and migrate on laminin A coated glass bottom dishes and observed by AiryScan confocal microscopy. Filmed at 1 frame/10 seconds, movie shows 7 frames/second.

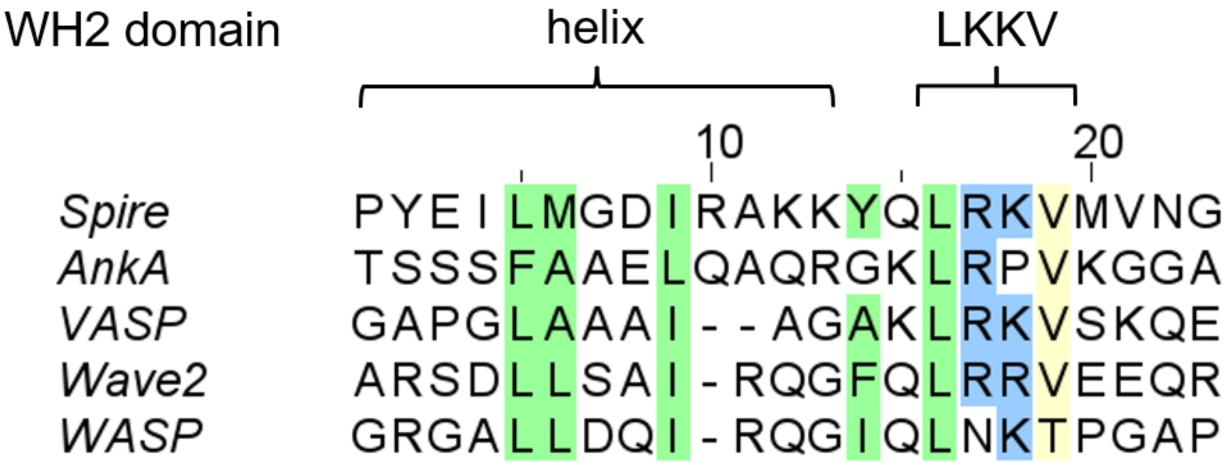


**Supplementary figure 1 Alignment of selected WH2 motifs with AnkA-ABM sequences**. The defining WH2 residues are highlighted green for hydrophobic, blue for basic, and yellow for small

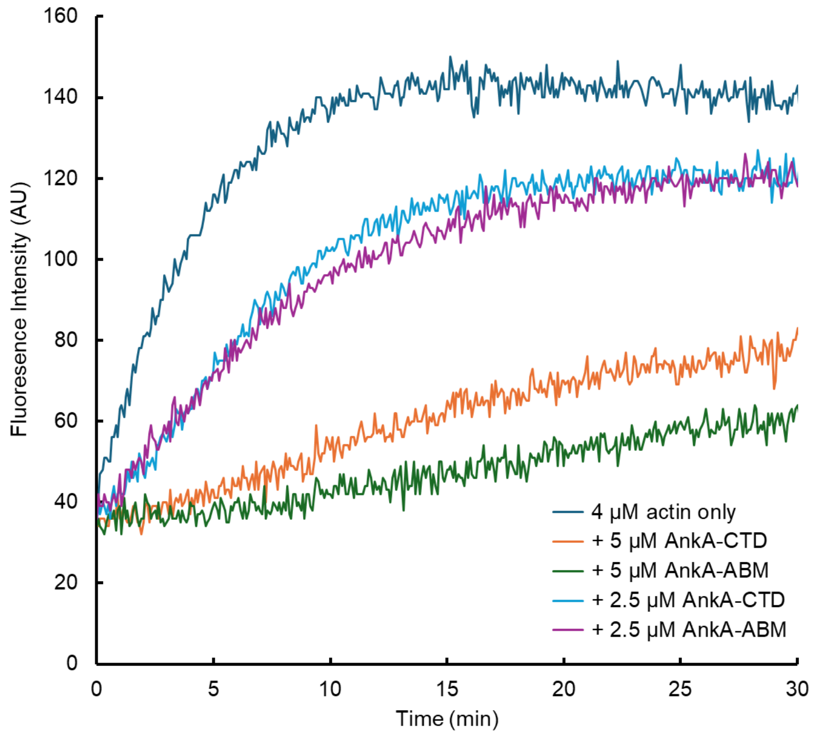


**Supplementary figure 2:** actin pyrene polymerisation assays in the presence of AnkA-CTD or AnkA-ABM peptide.

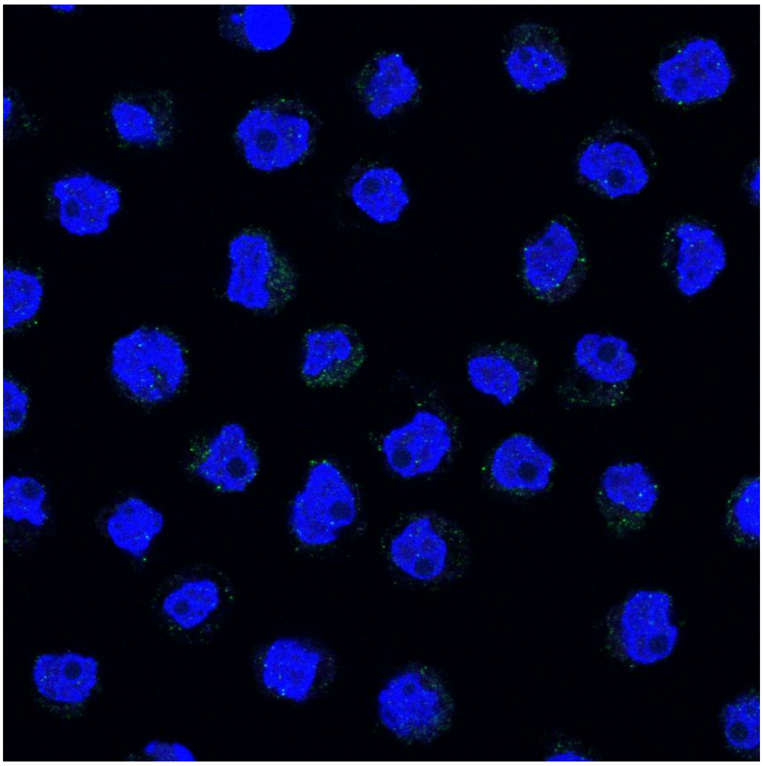

**Supplementary figure 3:** staining of uninfected HL-60 cells with anti-AnkA, anti-Abi1, and Hoechst 3332. Negligible staining of AnkA was detected whereas Abi1 has a generalized distribution in the cytoplasm.

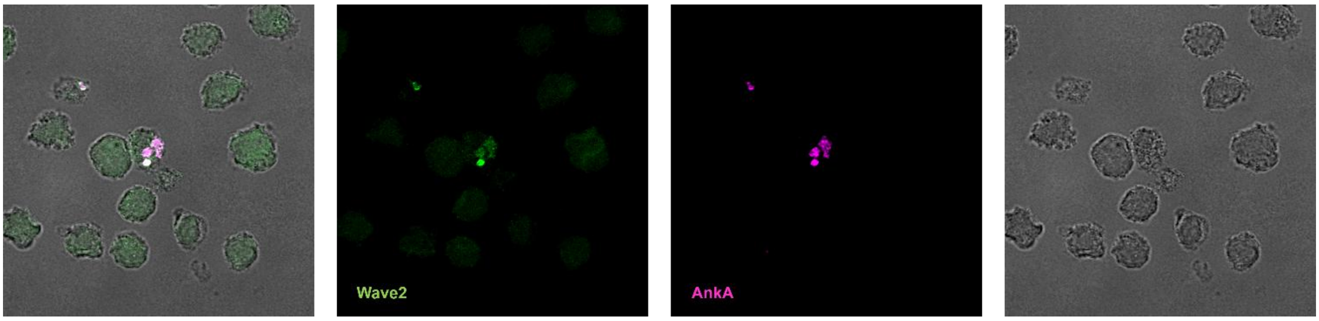


**Supplementary figure 4**. Confocal immunofluorescence microscopy images of *A. phagocytophilum* infected HL-60 cells stained with anti-AnkA and anti-Wave2.

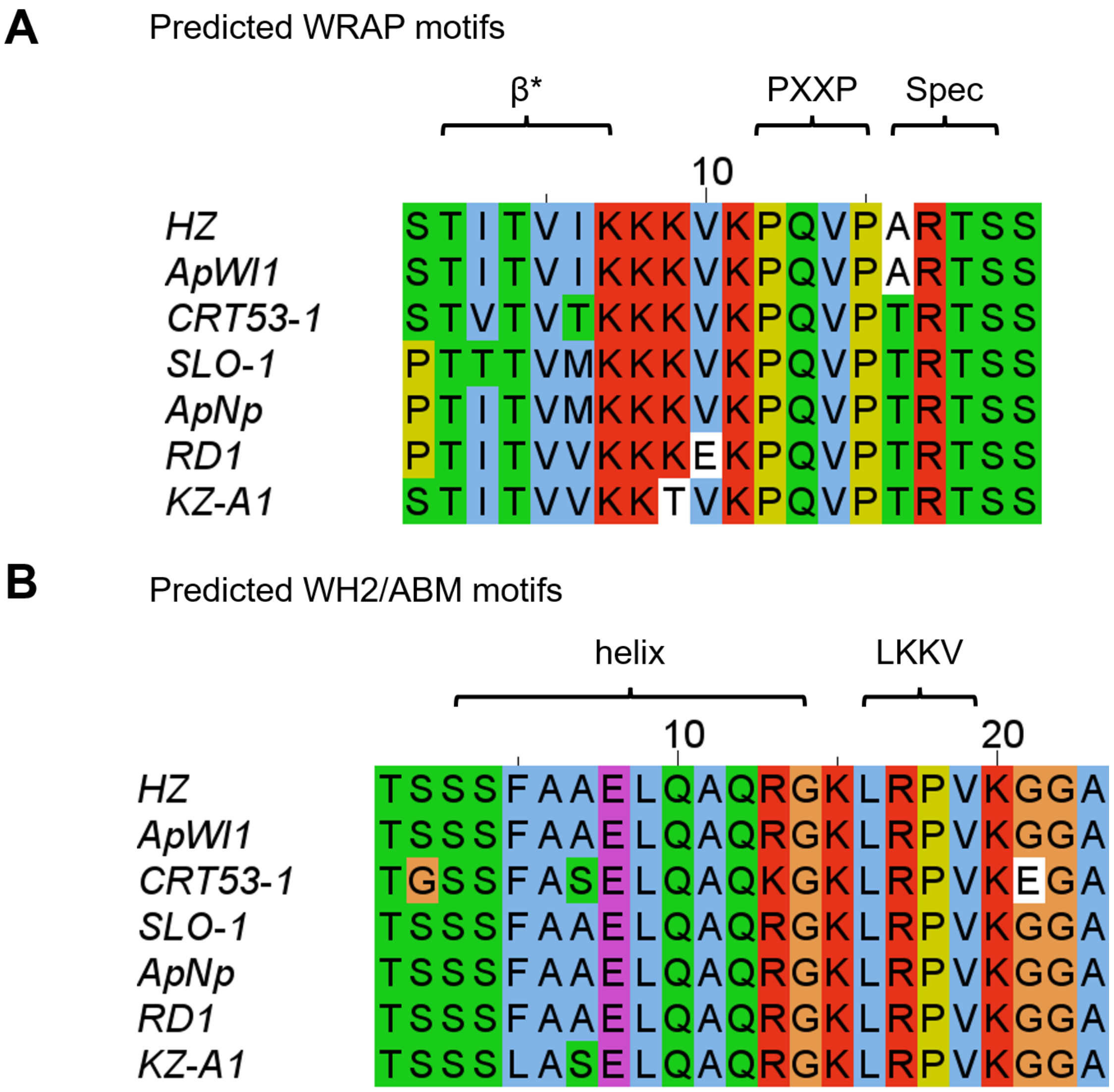


**Supplementary figure 5.** Alignment of A. WRAP and B. ABM motifs of AnkA from diverse *A. phagocytophilum* isolates. Sequences are from the following sources: HZ, *A. phagocytophilum* HZ, USA; ApWI1, *A. phagocytophilum* Wisconsin 1, USA; CRT53-1, *Anaplasma phagocytophilum* CRT53-1, USA; SLO-1, *A. phagocytophilum* from a human patient in Slovenia; ApNp, *A. phagocytophilum* from a horse in USA; RD1, *A. phagocytophilum* from a roe deer in Germany; KZ-A1, *A. phagocytophilum* from a human patient in South Korea.

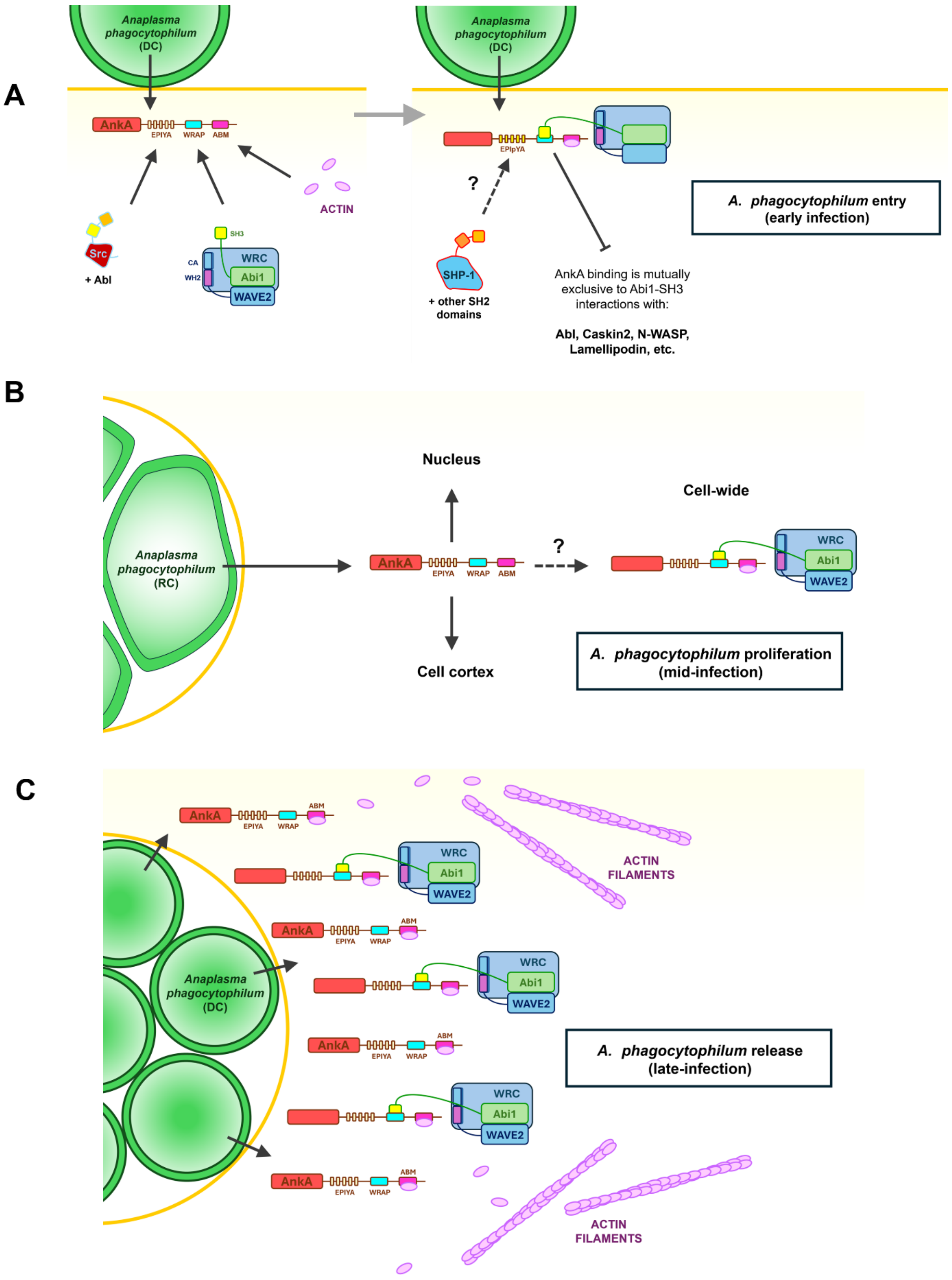
A
Anaplasma phagocytophilum (DC)
AnkA
EPIYA
WRAP
ABM
ACTIN
Src
+ Abl
SH3
CA
WH2
WRC
Abi1
WAVE2
EPIpYA
?
SHP-1
+ other SH2 domains
AnkA binding is mutually exclusive to Abi1-SH3 interactions with:
Abl, Caskin2, N-WASP, Lamellipodin, etc.
A. phagocytophilum entry (early infection)
B
Anaplasma phagocytophilum (RC)
Nucleus
Cell-wide
Cell cortex
A. phagocytophilum proliferation (mid-infection)
C
ACTIN FILAMENTS
A. phagocytophilum release (late-infection)

**Supplementary figure 6.** Model of AnkA functions during *A. phagocytophilum* infection of vertebrate host cells. A. During early infection AnkA is secreted into host cells and tyrosine phosphorylated on EPIYA-like motifs and binds to host actin and the WRC. Binding of the Abi1-SH3 domain of the WRC by AnkA is mutually exclusive with its interaction with other SH3-binding proteins. AnkA has a putative role in either facilitating WRC-dependent entry or influencing signalling pathways during the biogenesis of the ApV. B. During bacterial proliferation, AnkA is not associated with the ApV and is instead found throughout the cell. During this stage AnkA might fulfil other roles at the cell cortex and nucleus as proposed elsewhere (see main text). It is not clear if AnkA exerts influence in the WRC at this stage, since it was not possible to discern whether AnkA and Abi1 co-localise diffusely across the cell. C. Late in infection, AnkA and the WRC cluster at the ApV in an absence of obvious F-actin structures. At this stage we propose that AnkA sequesters cellular WRC, thereby influencing WRC-driven processes (e.g. cell migration, phagocytosis, etc), and (due to the high density of AnkA at the ApV) inhibits actin polymerisation local to the inclusion prior to exocytosis or host cell lysis.

**Supplementary table 1: Data collection and refinement statistics.**

| | **Abi1-SH3:AnkA-WRAP** |
|---|---|
| **Wavelength** | 0.9999 Å |
| **Resolution range** | 32.37 - 1.69 (1.73 - 1.69) |
| **Space group** | C 1 2 1 |
| **Unit cell** | 98.89 34.62 64.83 90 128.01 90 |
| **Total reflections** | 139182 (199) |
| **Unique reflections** | 25489 (192) |
| **Multiplicity** | 5.5 (1.0) |
| **Completeness (%)** | 97.76 (82.63) |
| **Mean I/sigma(I)** | 15.52 (2.36) |
| **Wilson B-factor** | 16.06 |
| **R-merge** | 0.0645 (0.3719) |
| **R-meas** | 0.07051 (0.5259) |
| **R-pim** | 0.028 (0.3719) |
| **CC1/2** | 0.996 (0.961) |
| **CC*** | 0.999 (0.99) |
| **Reflections used in refinement** | 19238 (1156) |
| **Reflections used for R-free** | 1924 (113) |
| **R-work** | 0.1616 (0.1950) |
| **R-free** | 0.2074 (0.2657) |
| **Number of non-hydrogen atoms** | 1778 |
| **macromolecules** | 1565 |
| **ligands** | 0 |

| | |
|---|---|
| **solvent** | 213 |
| **Protein residues** | 198 |
| **RMS(bonds)** | 0.010 |
| **RMS(angles)** | 1.08 |
| **Ramachandran favored (%)** | 97.37 |
| **Ramachandran allowed (%)** | 2.63 |
| **Ramachandran outliers (%)** | 0.00 |
| **Rotamer outliers (%)** | 0.00 |
| **Clashscore** | 1.29 |
| **Average B-factor** | 20.83 |
| **macromolecules** | 19.52 |
| **solvent** | 30.42 |

Statistics for the highest-resolution shell are shown in parentheses.

**Supplementary table 2: Data collection and refinement statistics.**

| | Actin:AnkA-ABM |
|---|---|
| **Wavelength** | 0.9000 |
| **Resolution range** | 65.5 - 2.84 (2.91 - 2.84) |
| **Space group** | P 43 21 2 |
| **Unit cell** | 131.01 131.01 206.4 90 90 90 |
| **Total reflections** | 1155916 (77702) |
| **Unique reflections** | 43093 (2821) |
| **Multiplicity** | 26.8 (27.5) |
| **Completeness (%)** | 99.59 (97.98) |
| **Mean I/sigma(I)** | 7.02 (0.50) |
| **Wilson B-factor** | 80.91 |
| **R-merge** | 0.5763 (9.675) |
| **R-meas** | 0.5874 (9.855) |
| **R-pim** | 0.1128 (1.865) |
| **CC1/2** | 0.996 (0.368) |
| **CC*** | 0.999 (0.733) |
| **Reflections used in refinement** | 42941 (2764) |
| **Reflections used for R-free** | 2162 (138) |
| **R-work** | 0.2507 (0.4309) |
| **R-free** | 0.2828 (0.4479) |
| **Number of non-hydrogen atoms** | 8704 |
| **macromolecules** | 8515 |

| | |
|---|---|
| **ligands** | 189 |
| **solvent** | 0 |
| **Protein residues** | 1094 |
| **RMS(bonds)** | 0.003 |
| **RMS(angles)** | 0.67 |
| **Ramachandran favored (%)** | 96.75 |
| **Ramachandran allowed (%)** | 2.97 |
| **Ramachandran outliers (%)** | 0.28 |
| **Rotamer outliers (%)** | 0.33 |
| **Clashscore** | 7.80 |
| **Average B-factor** | 101.43 |
| **macromolecules** | 101.74 |
| **ligands** | 87.29 |
